# A novel pipeline for the validation of manganese chelators for the treatment of manganese overload

**DOI:** 10.64898/2026.05.12.724311

**Authors:** Hendrik Vogt, Chiara Pojani, Jack Devonport, Andrew McGown, George Firth, Ivan Doykov, Valeria Nikolaenko, Sofia Anagianni, Leonardo E Valdivia, Youssef Khalil, Nikolett Bodnár, Csilla Kállay, Chris Dadswell, Ramon Gonzalez-Mendez, Rupert Purchase, Frances M. Platt, Flavia C. M. Zacconi, Amy F. Geard, Wendy E. Heywood, Kevin Mills, Philippa B. Mills, Ahad A. Rahim, Jason Rihel, Stephen W. Wilson, George E. Kostakis, John Spencer, Karin Tuschl

**Affiliations:** UCL GOS Institute of Child Health, University College London, London, UK; Department of Chemistry, School of Life Sciences, University of Sussex, Brighton, UK; School of Biomedical Engineering & Imaging Sciences, King’s College London, St. Thomas’ Hospital, London, UK; Department of Cell and Developmental Biology, University College London, London, UK; Center for Integrative Biology, Facultad de Ciencias, Ingeniería y Tecnología, Universidad Mayor, Santiago, Chile; Escuela de Biotecnología, Facultad de Ciencias, Ingeniería y Tecnología, Universidad Mayor, Santiago, Chile; Department of Inorganic and Analytical Chemistry, University of Debrecen, Debrecen, Hungary; Department of Pharmacology, University of Oxford, Oxford, UK; Department of Organic Chemistry, Faculty of Chemistry and of Pharmacy, Pontificia Universidad Católica de Chile, 4860 Av. Vicuña Mackenna, 7820436 Macul, Chile; Institute for Biological and Medical Engineering, Schools of Engineering, Medicine and Biological Sciences, Pontificia Universidad Católica de Chile, Macul, Chile; School of Pharmacy, University College London, London, UK; Department of Analytical, Environmental and Forensic Sciences, School of Cancer and Pharmaceutical Sciences, King’s College London, London, UK

## Abstract

Manganese neurotoxicity, arising from environmental overexposure or inherited transporter disorders due to pathogenic variants in SLC30A10 and SLC39A14, leads to manganism, a debilitating Parkinsonian movement disorder. Alhtough chelation therapy can partially reverse neuropathology, current clinical practice relies on intravenous CaNa2EDTA, which is burdensome and poorly suited for long-term use. Consequently, there remains a significant unmet need for more effective, orally bioavailable chelators. This study aimed to establish and validate a pipeline for identifying and assessing novel ligands that attenuate manganese neurotoxicity and support preclinical translational development.

Based on the structural features of manganese-based MRI contrast agents, we selected two chelators, N-picolyl-N,N′,N′-trans-1,2-cyclohexylenediaminetriacetic acid (H3PyC3A) and ethylenediaminetetraacetic acid−benzothiazole aniline (H4EDTA-BTA), and their methyl ester derivatives, Me3PyC3A and Me4EDTA-BTA. These were evaluated in vivo using zebrafish (slc39a14U801/U801) and mouse (Slc30a10KO/KO) models of manganese overload.

H3PyC3A and Me3PyC3A demonstrated greater manganese-mobilizing efficacy than CaNa2EDTA, improving locomotor behavior in slc39a14U801/U801 zebrafish. In Slc30a10KO/KO mice, intravenous administration confirmed selective in vivo chelation of excess manganese over physiological concentrations of zinc and copper. Although oral bioavailability was low (<1%), long-term oral administration of H3PyC3A modestly reduced liver and brain Mn accumulation, suggesting an added benefit of oral administration via gastrointestinal chelation.

This integrated in vitro to in vivo pipeline provides a robust and scaleable approach for the development of next-generation Mn chelators. Slc39a14U801 loss-of-function zebrafish enable high throughput identification of candidate compounds while Slc30a10KO/KO mice offer a clinically relevant disease model for pharmacokinetic profiling and proof-of-concept validation.

## Introduction

Manganese (Mn) is an essential trace metal that is critical for brain physiology and development (1, 2). As a constituent of metalloenzymes, Mn is involved in neurotransmitter recycling, ammonia and energy metabolism, glycosylation, and mitochondrial antioxidant defence (3, 4). Homeostasis of Mn levels in the body is tightly regulated through intestinal absorption and hepatobiliary secretion of Mn into the gastrointestinal tract (4–7). In human tissues, Mn exists primarily as divalent Mn(II), with smaller amounts present as trivalent Mn(III) ions (8). Increased levels of Mn are neurotoxic, leading to “manganism”, which is characterized by dystonia-parkinsonism, psychiatric and intellectual disability (4). Excess Mn accumulates in the basal ganglia, particularly the globus pallidus, where it causes distinct brain magnetic resonance imaging (MRI) appearances with pronounced hyperintensity of the globus pallidus on T1-weighted imaging (9).

Acquired Mn overload occurs in environmental and occupational overexposure as well as end-stage liver disease due to impaired biliary Mn excretion (5, 6). Recently, inherited forms of manganism have been identified that are caused by pathogenic variants in the Mn transporter encoding genes *SLC30A10* and *SLC39A14*, named hypermanganesemia with dystonia (HMNDYT) 1 and 2, respectively (10–14). In HMNDYT1, in addition to the neuropathology, hepatic Mn deposition leads to liver disease, polycythemia and depletion of iron stores (10).

There is also extensive evidence that metal dyshomeostasis plays a role in common neurodegenerative disorders, suggesting that targeting metal imbalance may be a relevant therapeutic approach (1, 15, 16). For instance, neuropathological features of Mn neurotoxicity, including oxidative stress, mitochondrial dysfunction, and impaired autophagy, are shared with neurodegenerative disorders such as Parkinson’s disease (PD) and Alzheimer’s disease (AD) (17). Manganism closely resembles PD, and inherited forms of PD due to pathogenic variants in *Parkin* and *ATP13A2* are associated with increased susceptibility to Mn toxicity (18, 19). Therefore, further investigation into the use of selective ligands towards Mn as chelation therapy is of paramount importance (15, 20).

In patients with inherited Mn transporter defects, intravenous CaNa_2_EDTA therapy can improve dystonia by increasing urinary Mn excretion and reducing Mn deposition in the brain and liver (8, 10, 12, 14, 21–24). However, in most cases some neurological sequelae persist with deterioration of motor function over time, possibly due to the limited pharmacokinetic properties of CaNa_2_EDTA leading to incomplete mobilization of Mn from the brain (11, 22, 25). Due to its hydrophilicity, CaNa_2_EDTA is distributed mainly in extracellular fluids and does not cross the cell membrane or blood-brain barrier (26, 27). In addition, the need for intravenous administration over multiple days every month causes hospital dependence and high socioeconomic costs (11, 25). Lack of metal selectivity can lead to adverse effects such as zinc (Zn) deficiency requiring Zn supplementation. Therefore, the development of an orally bioavailable chelator with selectivity towards Mn and optimized pharmacokinetic properties remains an unmet clinical need with the potential to substantially improve quality of life for patients affected by Mn-associated neurodegenerative diseases.

A variety of ligands that bind to Mn have found uses in medicine (8). For example, Mn complexes are used as superoxide dismutase mimics in model systems and non-toxic alternatives to gadolinium (Gd)-based magnetic resonance imaging (MRI) contrast agents (28–31), which are rapidly eliminated *in vivo* (32, 33). One such contrast agent is the complex Mn-PyC3A (34), derived from its ligand H_3_PyC3A (N-picolyl-N,N′,N′-trans-1,2-cyclohexylenediaminetriacetic acid), which was designed to replace Gd-based contrast agents with similar relaxivity and pharmacokinetics. Mn-EDTA-BTA, derived from its ligand H_4_EDTA-BTA (ethylenediaminetetraacetic acid−benzothiazole aniline), was developed for use as a liver-specific MRI contrast agent with high complex stability (35). This complex is taken up by hepatocytes and eliminated via hepatobiliary and renal pathways.

We hypothesized that the ligands H_3_PyC3A and EDTA-BTA could be repurposed as Mn selective chelators capable of reducing excess Mn levels *in vivo* (34–36). Previous studies suggest that their corresponding Mn complexes, Mn-PyC3A and Mn-EDTA-BTA, are highly resistant to transmetallation (metal exchange) indicating that *in vivo* displacement and redistribution of metal ions are unlikely, a prerequisite for chelator development (35, 36). To enhance pharmacokinetic properties, we also synthesized the more lipophilic methyl ester prodrugs Me_3_PyC3A and Me_4_EDTA-BTA (37), designed to improve membrane permeability and oral bioavailability, since both H_3_PyC3A and H_4_EDTA-BTA are hydrophilic with calculated LogP_o/w_ values of -0.55 and 0.16, respectively (38–40). To test the efficacy and clinical potential of repurposed Mn chelators we used two complementary animal models of Mn accumulation, *slc39a14* loss-of-function zebrafish that allow high throughput screening of candidate compounds, and *Slc30a10* knockout mice to assess pharmacokinetic behavior and establish proof-of-concept. Our repurposing strategy together with a preclinical translational validation pipeline provide an ideal framework for further drug development.

## Methods

### Chelator synthesis

Solvents and reagents were purchased from commercial suppliers and used without further purification. Chemical shifts are reported in parts per million (ppm), referenced to residual solvent peak. 1H, 13C NMR spectroscopy was carried out on Varian 500 or 600 MHz spectrometers. Liquid chromatography-mass spectrometry (LC-MS) measurements were performed on a Shimadzu LCMS-2020 using a Gemini® 5µm C18 110 Å column. % Purity measurements were performed using a 30-minute run (H_2_O, MeCN, 0.1% HCO_2_H) (5 min at 5%, 5–95% over 20 min, 5 min at 95%). UV detection was carried out at 254 nm. High-resolution accurate mass spectrometry measurements were obtained using a Waters Xevo G2 Q-ToF HRMS equipped with an ESI source and MassLynx software. ESI source parameters were as follows: capillary voltage 3.0 kV, sampling cone 35au, extraction cone 4au, source temperature 120 °C and desolvation gas 450 °C with a desolvation gas flow of 650 L/h and no cone gas. Mass spectra were acquired in resolution mode over an m/z range of 100 and 1500 Da. Additionally, a Waters (Wilmslow, Cheshire, UK) Acquity H-Class UHPLC chromatography system with column oven was used, connected to a Waters Synapt G2 HDMS high-resolution mass spectrometer. Chromatographic purification was carried out on an ISCO purification unit using Redisep silica gel columns. For the synthesis of H_3_PyC3A, Me_3_PyC3A, H_4_EDTA-BTA and Me_4_EDTA-BTA see **Figures S1-16**.

### Potentiometric studies of H_3_PyC3A in aqueous solution

The protonation constants of H_3_PyC3A and the stability constants of the Mn(II) and Zn(II) complexes were determined by pH-metric titration in 4mL samples at 1mM concentration. The metal ion to ligand ratio was 1:1. A MOLSPIN pH-meter equipped with a 6.0234.100 combined glass electrode (Metrohm) was used for pH measurements (in the pH range 2.7–11.5), while the dosing of the titrant was made with a MOL-ACS microburette. During the measurements argon was bubbled through the samples to ensure the absence of oxygen and carbon dioxide. All pH-potentiometric measurements were carried out at a constant ionic strength of 0.2 M KCl and at a constant temperature (298 K). Protonation constants and overall stability constants (log β) were calculated by means of the computational programs, PSEQUAD and SUPERQUAD.

### Potentiometric studies of H_3_PyC3A and H_4_EDTA-BTA in DMSO:water

The titrations were performed with a carbonate free stock solution of potassium hydroxide (prepared in 30:70 (w/w%) DMSO-water mixture) of known concentration. Mettler-Toledo T50 pH-meter equipped with a Metrohm glass electrode (model 6.0234.100) was used for pH measurements. During the measurements argon was bubbled through the samples to ensure the absence of oxygen and carbon dioxide. All pH-potentiometric measurements were carried out at a constant ionic strength of 0.2 M KCl and at a constant temperature (298 K). Protonation constants and overall stability constants (log β) were calculated by means of the computational programs PSEQUAD and SUPERQUAD.

### Sex as a biological variable

In line with the latest guidelines on the inclusion of sex as a biological variable (SABV) in preclinical studies, both male and female animals were included in the studies.

### Zebrafish strains and husbandry

Zebrafish (*Danio rerio*) were raised under standard conditions at 28 °C. The *slc39a14^U801^* line was available from previous studies (14). Ethical approval for zebrafish experiments was obtained from the Home Office UK under the Animal Scientific Procedures Act 1986.

### Locomotor zebrafish behavior assay

The behavioral assay was conducted as described previously (14). Briefly, zebrafish embryos and larvae were raised on a 14 h:10 h light:dark cycle. Single larvae were transferred to each well of a flat-bottom, clear polystyrene 96 well plate (Whatman) in fishwater (650 µL) at 4 days post fertilization (dpf). The 96-well plate was maintained at a constant temperature (26 °C) and exposed to a 14 h:10 h white light:dark schedule with constant infrared illumination within a custom-modified Zebrabox (Viewpoint LifeSciences). MnCl_2_ and ligands were added directly to the fishwater from 2 dpf. The locomotor behavior of zebrafish larvae was tracked from 4 to 7 dpf using an automated video tracking system (Viewpoint LifeSciences). Larval movement was recorded using Videotrack Quantization mode. The Videotrack detection parameters were empirically defined for clean detection of larval movement with minimal noise. A custom-designed Matlab code was used to extract the average activity data of each larva (14).

### Cardiac injection of ligands

Zebrafish larvae raised in 50µM MnCl_2_ from 2 to 6 dpf were injected directly into the heart at 4 and 5 dpf with 100 pg – 5 ng of ligand (H_3_PyC3A, CaNa_2_EDTA, or Me_3_PyC3A) after immobilization in 1.5% low melting point agarose (prepared in fishwater containing 0.02% MS-222) using a pressure-pulsed Picospritzer II (General Valve Corp.). Following injection, the larvae were removed from the agarose and maintained in fishwater until analysis. ICP-MS analysis was performed at 6 dpf.

### Mouse strains and husbandry

All animal experiments were performed in accordance with the Animal Scientific Procedures Act 1986. SLC30A10^KO/KO^ mice (*Mus musculus*) in C57BL/6N background were rederived from sperm, kindly provided by Prof Bartnikas (Brown University) (41). Mice were maintained on a 12 h light:12 h dark cycle with controlled temperature and humidity. Ear biopsies were collected at post-natal day 14 for genotyping, after which mice were weaned at post-natal day 21. Upon weaning, mice were group-housed in individually ventilated cages. Animals were randomly allocated to each experimental group. Animals had ad libitum access to standard chow (Teklad Global 18% Protein Rodent Diet 2018) and water throughout the experiments.

### Genotyping

DNA was extracted from ear biopsies with 50 μL extraction solution (Sigma E7526) and incubation at 95 °C for 15 minutes followed by addition of 50 μL neutralization solution (Sigma N3910). Allele specific primer touchdown PCR with betaine at a final concentration of 0.5 M was used to identify the genotype of each animal using the following primers: WT_Fw 5’ TCCGTGGTTGTGGTCATCA 3’, WT_Rv 5’ GTGTTTGTGGTCCTTGCTCA 3’, Mut_Fw 5’ AGGCCATCTTTCAGGAGTCA 3’, and Mut_Rv 5’ ATGAACTGATGGCGAGCTCA 3’.

### Chelator administration

#### IV and IP administration

H_3_PyC3A (20 mg/kg body weight) was dissolved in PBS, filter sterilised and administered via tail vein or intraperitoneal route.

#### Oral administration

To optimize stability and solubility, the HCl-salt of PyC3A (HCl-PyC3A) was used for all experiments involving oral administration. In brief, PyC3A was dissolved in minimal 1,4-dioxane with stirring. The resulting solution was then treated with 4N HCl in 1,4-dioxane (3.0 eq) and stirred at RT overnight. Upon completion the reaction mixture was concentrated to residue and resuspended in minimal 1,4-dioxane before being dried by lyophilization over night to yield a flocculant, colorless solid in quantitative yields (**Figure S1**).

For single oral administration, HCl-PyC3A (80 mg/kg body weight) was dissolved in PBS, filtered sterilized and administered by means of a micropipette.

For long-term studies, HCl-PyC3A (0.8 mg/mL) was dissolved in 100 mL of milli-Q water and provided *ad libitum* from weaning up to 10 weeks of age.

### Tissue collection

Mice were euthanized by transcardial perfusion with 10 mL of cold PBS under terminal isoflurane anesthesia, and brains, major visceral organs, and blood samples collected. Tissue samples were snap frozen on dry ice and stored at -80 °C until analysis.

### Blood sampling and hematology

Whole blood was collected on blood spot cards for LC-MS/MS analysis, in K_2_EDTA-coated collection tubes (BD Microtainer) for the determination of full blood counts or in Eppendorf tubes for ICP-MS measurements. Blood was sampled via tail vein bleeding from live mice or cardiac puncture under terminal anesthesia. The number of red blood cells was determined using the Sysmex XE-5000™ Automated Hematology System (Sysmex Corporation, Japan).

### LC-MS/MS assay of PyC3A–metal complexes from dried blood spot samples

PyC3A–metal complexes were quantified from dried blood spot (DBS) samples using hydrophilic interaction liquid chromatography tandem mass spectrometry (HILIC–LC–MS/MS). Circular 3 mm DBS punches were extracted with 50µL of an acetonitrile–methanol–water mixture (50:30:20, v/v/v) containing 5mM ammonium acetate (pH 7.6). The samples were shaken for 30 min at room temperature, sonicated for 15 min and centrifuged at 16,000 × g for 10 min. An aliquot of the supernatant was transferred to a polypropylene vial for subsequent analysis.

Chromatographic separation was conducted on an ACQUITY Premier BEH Amide column (1.7 µm, 2.1 × 100 mm) maintained at 40 °C, with a flow rate of 0.2 mL/min and an injection volume of 3µL. The mobile phase composition and gradient conditions were optimized for the retention of hydrophilic PyC3A complexes with extended column equilibration and conditioning to ensure stable HILIC performance. Detection was performed on a triple quadrupole mass spectrometer Xevo TQ-S operated in the negative electrospray ionization mode. Multiple reaction-monitoring (MRM) transitions were employed for PyC3A complexes with Mn²⁺, Co²⁺, Cu²⁺ and Zn²⁺, as well as for free PyC3A (detailed in Supplementary Information). The source and collision cell parameters were optimized for each analyte. In the absence of suitable internal standards, data were median-normalized to account for run-to-run variability.

Full details of the reagents, chromatographic conditions, mass spectrometric settings, and MRM transitions are provided in the Supplementary Methods.

### Urine collection and creatinine assay

Urine samples were collected by aspirating urine from a sterile Petri dish after manually inducing urination through gentle pressure on the lower abdomen. Urine samples were stored at -80 °C until further processing. Urine creatinine levels were measured by LC-MS/MS using established methods (42).

### Inductively coupled plasma mass spectrometry (ICP-MS) analysis of metal ions

Mouse tissues were incubated in open lid 1.5 mL Eppendorf tubes in a hybridization oven at 75 °C and weighed before and after to determine the wet and dry weight, respectively. Mouse tissue samples or 10 zebrafish larvae were digested in 200 µL concentrated nitric acid at 95 °C until dry and resuspended in 1 mL of 3% nitric acid. 50 μL of whole blood or urine were added to 1.95 mL of 3% nitric acid (Fisher) and incubated at 85 °C for 4 h. After centrifugation, the supernatant was analyzed by ICP-MS. Metal concentrations were measured using an Agilent 7500ce ICP-MS instrument with collision cell (in He mode) and Integrated Autosampler (I-AS) using ^72^Ge as internal standard. The following experimental parameters were used: a) plasma: RF power 1500 W, sampling depth 8.5 mm, carrier gas 0.8 L/min, make-up gas 0.11 L/min; b) quadrupole: mass range 1-250 amu, dwell time 100 msec, replicates 3, integration time 0.1 sec/point. Calibration solutions were prepared for each element between 0 and 200 ng/mL using certified reference standards (Fisher Scientific, UK).

### Statistical analysis

Statistical analysis was performed using GraphPad Prism 10 Software. For comparison of two groups Student’s t-test was used. For comparisons with more than two conditions, one-way ANOVA with Tukey’s post hoc tests was used for normally distributed data and Kruskal-Wallis multiple analysis with Dunn’s multiple comparison’s test for data that did not follow a normal distribution.

### Study Approval

Ethical approval was obtained for all animal studies from the Home Office UK under the Animal Scientific Procedures Act 1986. Mouse experiments were performed in accordance with the ARRIVE Guidelines.

## Results

### The ligands H_3_PyC3A and H_4_EDTA-BTA form stable chelate complexes with Mn *in vitro*

We synthesized both ligands, H_3_PyC3A and H_4_EDTA-BTA (**Figure 1A**), as well as their methyl ester prodrugs. The reported protocol for synthesis of H_3_PyC3A was slightly modified by reducing the number of synthetic steps (**Figure S1**) (34, 36). H_4_EDTA-BTA was isolated in high purity according to a published protocol (35). In both cases, the corresponding methyl ester prodrugs (Me_3_PyC3A and Me_4_-EDTA-BTA) were synthesized using methyl bromoacetate in the exhaustive *N*-alkylation of the corresponding ligands with the aim of targeting endogenous esterases to slowly release the active (H_3_PyC3A and H_4_EDTA-BTA) ligands (**Figures S1-S18, Table S1**) (37, 43). Deprotonation and stability constants were determined by potentiometric studies (**Figures S19-21, Tables S2-S5**). The stability constants for the Mn(II) and Zn(II) complexes of both ligands and p[M] values directly compare the affinity of the ligands towards a metal center (**Figure 1B**). The studies of H_3_PyC3A in an aqueous system gave a value of pMn_7.4_ = 8.14, which is in good agreement with previous studies (pMn_7.4_ = 8.17) (36). However, we noted that H_3_PyC3A has an even higher affinity for Zn(II) ions under the same conditions with a pZn_7.4_ = 9.60. Upon altering the solvent system, DMSO:water (70:30), we identified that the pMn_7.4_ value decreased to pMn_7.4_ = 7.53 (**Figure 1B**). However, pMn_7.4_ remained an order of magnitude higher than that of H_4_EDTA-BTA in the same system, pMn_7.4_ = 6.45 (**Figure 1B**), suggesting a higher Mn binding affinity for H_3_PyC3A. Previous studies suggest that the formed Mn chelates, Mn-PyC3A and Mn-EDTA-BTA, are highly resistant to transmetallation (metal exchange) indicating that *in vivo* displacement and redistribution of metal ions are unlikely, a prerequisite for chelator development (35, 36).

**Figure 1.**
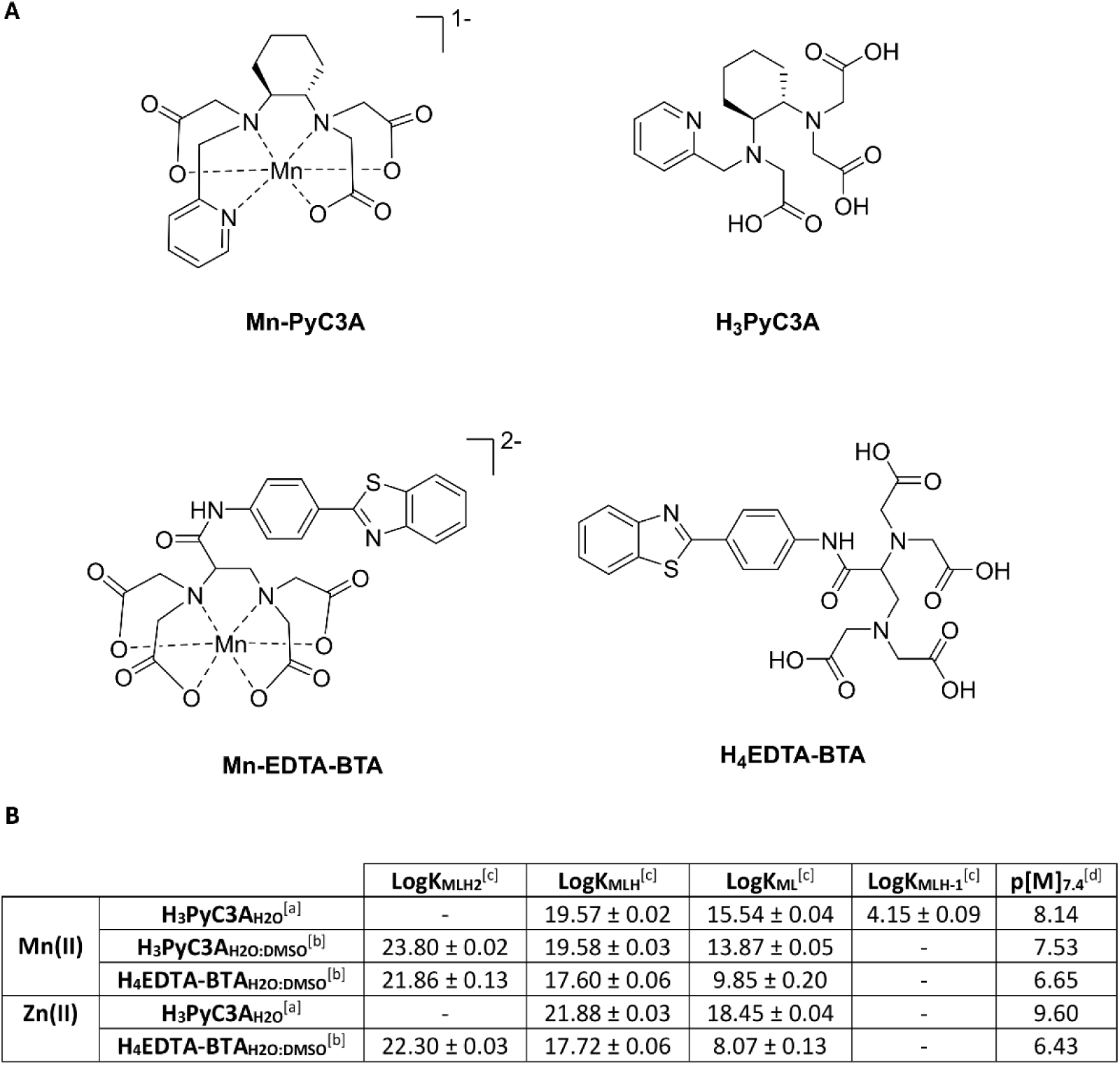
H_3_PyC3A and H_4_EDTA-BTA form stable chelate complexes with Mn *in vitro*. **A** Mn complexes Mn-PyC3A and Mn-EDTA-BTA assuming a 2+ oxidation state for Mn (left), and the corresponding organic scaffolds (right). **B** Stability constants of the ligands for their corresponding Mn(II) and Zn(II) complexes. [a]: Calculated from potentiometric data collected in an aqueous solution. [b]: Calculated from potentiometric data collected in a H_2_O:DMSO = 30:70 (w/w%) system. [c]: Defined as: KMLH_2_; [MLH_2_]/([MLH][H])], KMLH; [MLH]/([ML][H]), KML; [ML]/([M][L]), KMLH-1; [MLH-1]/([ML][H-1])]. [d]: p[M] = -log[M]free at pH = 7.4 when [M] = [L] = 10µM.

### *In vitro* pharmacokinetic studies validate H_3_PyC3A safety for *in vivo* application

*In vitro* pharmacokinetic studies on H_3_PyC3A, Me_3_PyC3A and H_4_EDTA-BTA (outsourced to Eurofins Scientific) confirmed low HepG2 cytotoxicity and excellent solubility for all three ligands (**Table 1**) (44, 45). H_3_PyC3A demonstrated high plasma stability and relatively low plasma protein binding. Me_3_PyC3A demonstrated rapid plasma activation; however, attempts to identify its metabolites were inconclusive, as only incomplete demethylation was observed, rather than the desired exhaustive conversion to PyC3A.

**Table 1.** Comparison of *in vitro* behavior of Me_3_PyC3A, H_3_PyC3A and H_4_EDTA-BTA. Nd, not determined. PB, plasma protein binding.

| Compound | HepG2<br>Cytotox %<br>inhibition<br>(100 μM)<br>MEAN | HepG2<br>Cytotox<br>EC50<br>(μM)<br>MEAN | Limit of<br>Solubility<br>pH 7 (μM)<br>MEAN | Plasma<br>Stability<br>(mouse)<br>T<sub>1/2</sub> (min)<br>MEAN | Plasma<br>Stability<br>(mouse) %<br>Remaining<br>After 2<br>Hours<br>MEAN | PPB<br>(mouse)<br>%<br>Unbound<br>MEAN | PPB<br>(mouse)<br>% Bound<br>MEAN |
| --- | --- | --- | --- | --- | --- | --- | --- |
| Me<sub>3</sub>PyC3A | 64.3 | 47.7 | >100 | 10.1 | 0 | nd | nd |
| H<sub>3</sub>PyC3A | 26.7 | >100 | >100 | >400 | 91 | 73.6 | 26.4 |
| H<sub>4</sub>EDTA-BTA | 32.3 | >100 | >100 | nd | nd | nd | nd |

### Treatment with H_3_PyC3A ameliorates Mn toxicity in zebrafish

*In vivo* efficacy of the above chelators relative to CaNa_2_EDTA was evaluated in *slc39a14^u801^*loss-of-function zebrafish (herein called *slc39a14^-/-^*), a model of the inherited Mn transporter defect HMNDYT2 (14). Similar to affected patients, upon MnCl_2_ exposure, *slc39a14^-/-^* larvae develop Mn deposition in the brain and impaired locomotor activity characterized by reduced average activity during the day and increased activity at night (46). Mutant larvae also display a lack of visual background adaptation, evident as darker pigmentation compared to wild-type zebrafish, which is associated with changes in visual behaviours (46). The small size of zebrafish larvae enables high-throughput screening, as their locomotor behavior can be tracked in a 96-well plate format (47).

The respective ligand was added to the fishwater from 2 days post fertilization (dpf), and larval movement was tracked continuously on a 14 hour day:10 hour night cycle from 4 to 7 dpf using an automated video tracking system as described previously (14). H_3_PyC3A treatment led to normalization of the average day and night activity of *slc39a14^-/-^*larvae upon MnCl_2_ treatment (**Figure 2A-B**), while CaNa_2_EDTA had minimal effects (**Figure 2C-D**), as observed previously (14). Total larval Mn levels, as determined by ICP-MS, almost normalized upon H_3_PyC3A treatment (**Figure 2E**, 95% reduction), while only some reduction was observed with CaNa_2_EDTA (**Figure 2F**, 29% reduction). Furthermore, H_3_PyC3A treated larvae normalized their pigmentation pattern while untreated and CaNa_2_EDTA treated larvae remained darkly pigmented with dispersion of their melanosomes (**Figures 2G-H**). H_4_EDTA-BTA on the other hand did not affect the motor activity of MnCl_2_ exposed *slc39a14^-/-^*larvae (**Figure S22**), and its methyl ester Me_4_-EDTA-BTA was toxic to zebrafish at 7 dpf. Hence, EDTA-BTA-based ligands were disregarded from further analysis.

**Figure 2.**
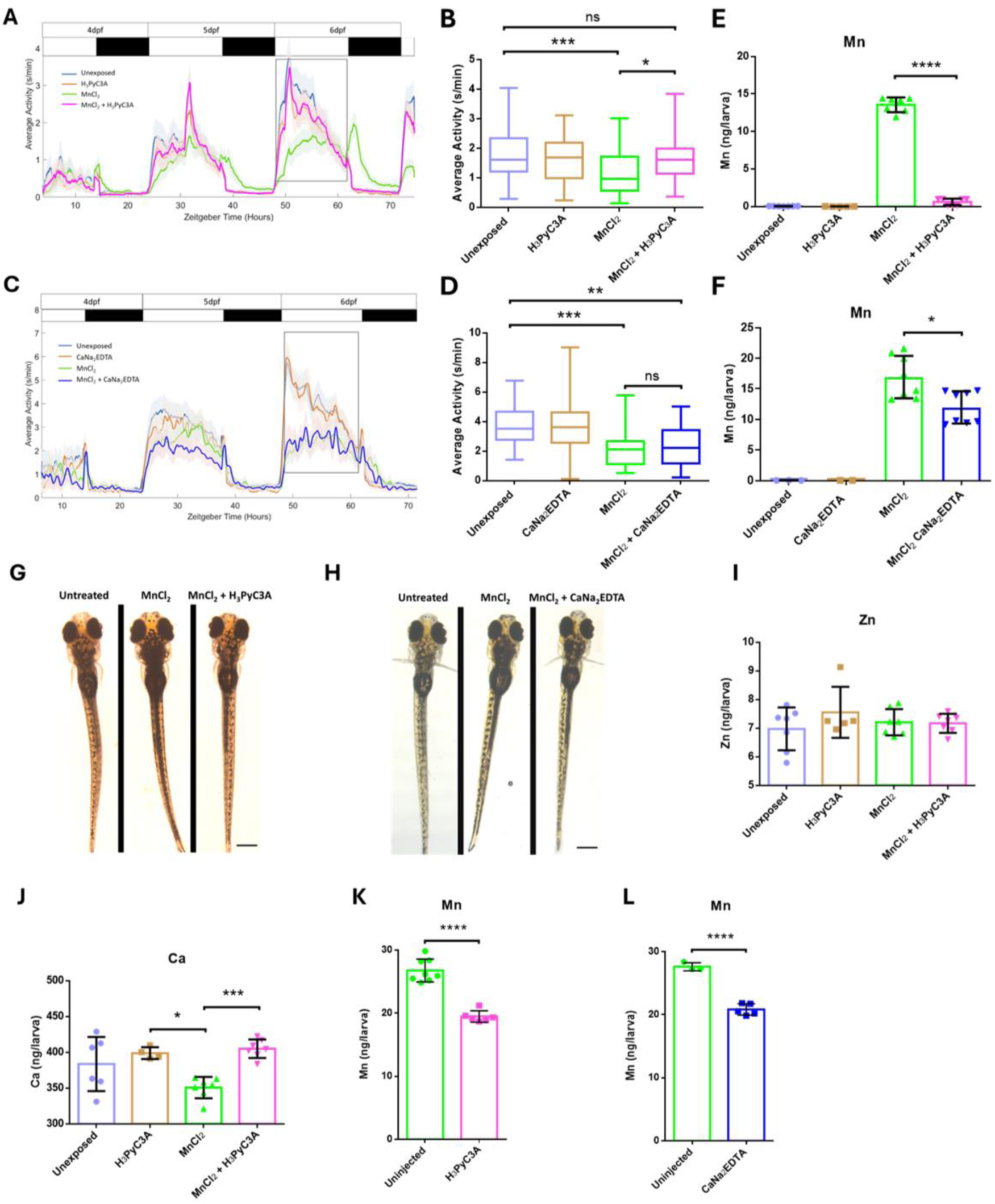
H_3_PyC3A treatment restores locomotion, pigmentation pattern and Mn levels in *slc39a14^-/-^* zebrafish. **A-B** Locomotor activity pattern during 4-6 dpf under **A** H_3_PyC3A (40 µM) and **B** CaNa_2_EDTA (40 µM) treatment from 2 dpf. Rectangles highlight the changes observed at 6 dpf. Boxplot (on right) showing average day activity at 6 dpf (n ≥ 24 per condition). Kruskal-Wallis multiple analysis with Dunn’s multiple comparison’s test; ns, not significant; A \*\*\**p* = 0.0005, \**p* = 0.0123. B \*\*\**p* = 0.0008, \*\**p* = 0.0044; **C-D** Representative brightfield images of the larval pigmentation patterns upon **C** H_3_PyC3A and **D** CaNa_2_EDTA treatment at 6 dpf. Size bar 300 µm. **E-F** Mn content of slc39a14^-/-^ larvae at 7 dpf determined by ICP-MS from pooled samples of 10 MnCl_2_ treated larvae (50 µM from 2 dpf) at 6 dpf following cardiac injection of 5ng **E** H_3_PyC3A or **F** CaNa_2_EDTA at 4 and 5 dpf. Student’s t-test. \*\*\*\**p* < 0.0001.

Given that the stability constant for the Mn(II) and Zn(II) complex suggested higher affinity of H_3_PyC3A to Zn(II) than Mn(II), we also determined the effects of H_3_PyC3A on larval levels of divalent metals other than Mn. However, no changes were observed for Zn (**Figure 2I**), suggesting an *in vivo* selectivity of H_3_PyC3A in conditions of Mn overload. We have previously shown that MnCl_2_ exposure lowers Ca levels in *slc39a14^-/-^* zebrafish (46). As expected, Ca levels normalized upon H_3_PyC3A treatment in addition to Mn levels (**Figure 2J**). To confirm that the observed reductions in Mn accumulation are due to the actions of H_3_PyC3A in larval zebrafish *in vivo* and not merely through chelation of Mn present in the fishwater, we injected H_3_PyC3A into the heart of MnCl_2_ treated *slc39a14^-/-^* zebrafish larvae and determined total Mn levels by ICP-MS. Cardiac injection of H_3_PyC3A led to a decrease in Mn levels by 27.2% compared to a 24.5% reduction by injection of CaNa_2_EDTA (**Figure 2K-L**). This confirms that H_3_PyC3A chelates Mn *in vivo* to a comparable extent as CaNa_2_EDTA when injected, while CaNa_2_EDTA has no effect by immersion. Similar to our observations during H_3_PyC3A immersion, H_3_PyC3A injection did not affect Zn levels while Fe levels were mildly reduced (**Figure S23**). Ca levels upon injection were unaffected, probably due to the less significant reduction in Mn levels compared to immersion in H_3_PyC3A (**Figure S23**).

Subsequently, we assessed the effect of the tri-methyl ester Me_3_PyC3A on locomotor activity. Similar to our observations with H_3_PyC3A, immersion in Me_3_PyC3A led to improvements of average day activity in MnCl_2_ treated *slc39a14^-/-^* larvae, and cardiac injection reduced larval Mn levels by 24% (**Figure S24**).

### Intravenous administration of H_3_PyC3A leads to Mn complex formation in a genotype dependent manner

Since H_3_PyC3A demonstrated promising Mn chelation potential in zebrafish, we further examined the pharmacokinetic and chelation behavior in a mouse model of HMNDYT1, a hereditary movement disorder characterized by systemic Mn overload (6–8). This mouse model carries a loss-of-function mutation in the efflux transporter SLC30A10, recapitulating key features of the human clinical phenotype including brain and liver Mn accumulation and polycythemia. First, we assessed the *in vivo* selectivity of H_3_PyC3A for Mn relative to other divalent metal ions following intravenous (iv) injection of H_3_PyC3A in *Slc30a10^-/-^* or wild-type (WT) mice. To quantify metal-H_3_PyC3A complex formation we developed an LC-MS/MS assay in dried blood spots in order to detect Mn-PyC3A, Zn-PyC3A and Cu-PyC3A (**Figure 3A**). After injection of H_3_PyC3A (20 mg/kg) we were able to detect the formation of all three metal complexes - Mn-PyC3A, Zn-PyC3A and Cu-PyC3A in both genotypes (**Figure 3B, S25A,B**). In *Slc30a10^-/-^* mice, the proportion of Mn-PyC3A was considerably higher compared to WT mice suggesting selective chelation of excess Mn in *Slc30a10^-/^*^-^ mice, consistent with the genotype-dependent Mn burden (**Figure 3B-C**). Formation of both, Zn-PyC3A and Cu-PyC3A did not differ significantly between genotypes (**Figure S25A,B**). We further confirmed that urinary Mn excretion increased upon administration of H_3_PyC3A which is renally excreted (34). Urinary Mn levels, determined by ICP-MS, increased significantly 30 minutes after iv injection of H_3_PyC3A, similar to observations with CaNa_2_EDTA (**Figure 3D-E**).

**Figure 3.**
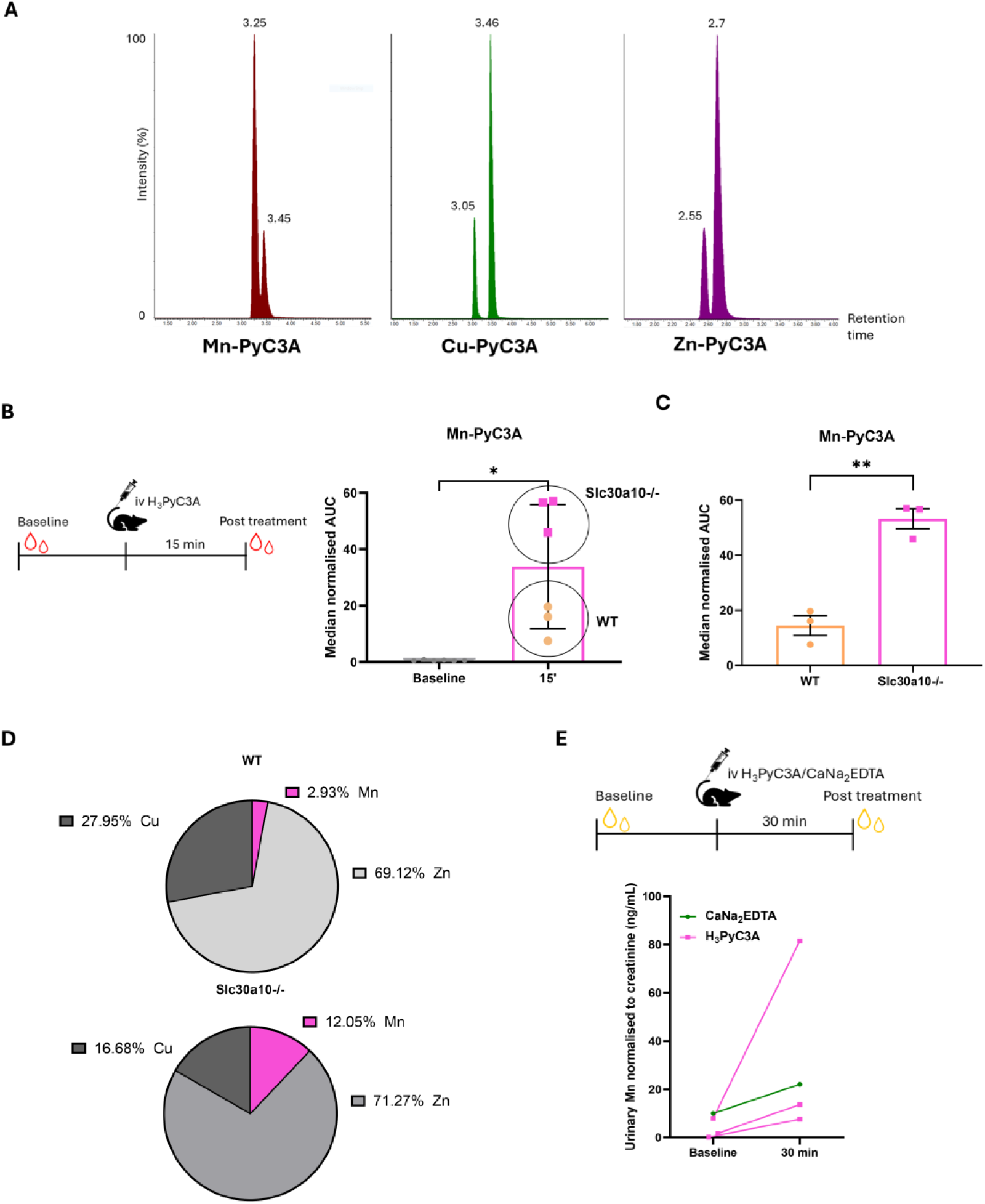
Intravenous administration of H_3_PyC3A leads to Mn complex formation in proportion to systemic Mn load. **A** Chromatograms showing the metal complexes Mn-PyC3A, Cu-PyC3A and Zn-PyC3A. Separation of the metal complexes was achieved using Hydrophilic Interaction Liquid Chromatography (HILIC) with an amide stationary phase. **B** Mn-PyC3A complex formation 15 minutes after iv injection of 20 mg/kg H_3_PyC3A in Slc30a10^-/-^ and WT mice (n = 3 per genotype). Data plotted as mean ± SD. One-way ANOVA, *p* < 0.0001. Tukey’s multiple comparisons test \*\**p* = 0.0031, ****p < 0.0001. **C** Pie chart demonstrating the fraction of Mn-PyC3A complex formation compared to Zn-PyC3A and Cu-PyC3A in Slc30a10^-/-^ and WT mice (n = 3 per genotype). **D** Urinary Mn concentration determined by ICP-MS 30 minutes following iv injection of either H_3_PyC3A or CaNa_2_EDTA (both at 20 mg/kg). Mn concentration was normalised to urinary creatinine to account for urine dilution^43^.

Overall, these results confirm the potential of H_3_PyC3A to reduce the systemic Mn load through selective Mn chelation and urinary excretion.

### H_3_PyC3A demonstrates poor oral bioavailability, but long-term oral administration may improve features of Mn overload

Current clinical interventions for Mn overload are limited to intravenous administration of CaNa_2_EDTA, which is associated with significant adverse effects (6–8, 18–20). To address this limitation, we explored the oral bioavailability and long-term therapeutic potential of H_3_PyC3A. For oral administration we used the HCl form of PyC3A for better stability and solubility. *Slc30a10^-/-^* mice received HCl-PyC3A via micropipette-guided oral administration, and whole blood was sampled 15 and 30 minutes post-treatment to determine metal-chelate formation over time by LC-MS/MS. We observed a modest increase in Mn-PyC3A complex concentration over 30 minutes, with a slight, gradual rise between 15 and 30 minutes following administration (**Figure 4A**) similar to observations with Cu and Zn-PyC3A concentrations (**Figure S25C,D**). Comparison of median normalized AUC values following intravenous (AUC ∼20–60; **Figure 3B**) versus oral administration (AUC ∼0.2–0.5; **Figure 4A**) indicates a low oral bioavailability of approximately 1%.

**Figure 4.**
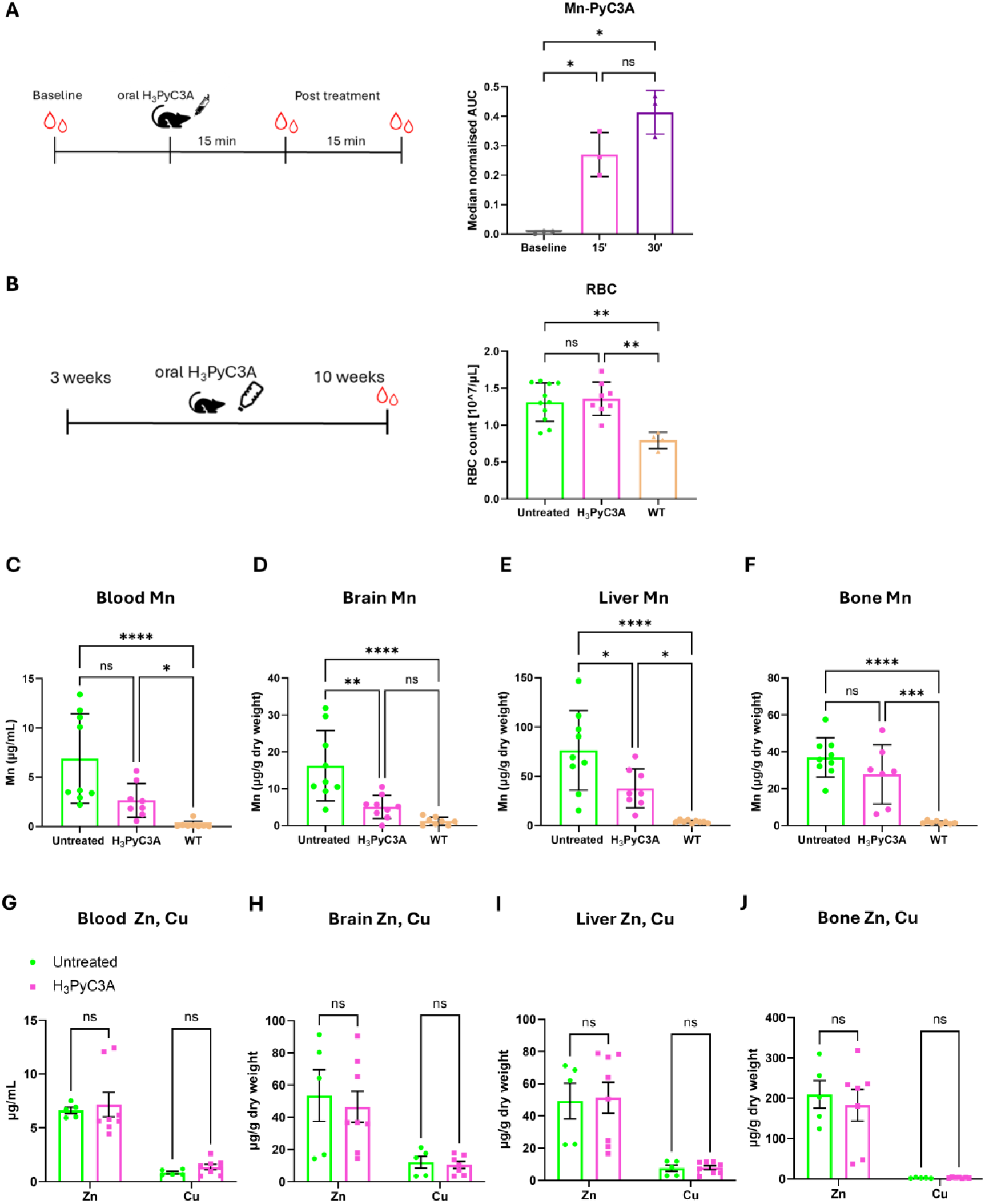
Oral administration of HCl-PyC3A reduces Mn accumulation despite poor oral bioavailability. **A** HCl-PyC3A (80 mg/kg) was administered orally to Slc30a10^-/-^ mice and Mn-PyC3A levels measured by LC-MS/MS (n = 3). Data plotted as mean ± SD. One-Way Anova (*p* = 0.0098) with Tukey’s multiple comparisons test: 15’ *p* = 0.0463, 30’ *p* = 0.0170. **B-F** Slc30a10^-/-^ mice were fed Milli-Q water supplemented with HCl-PyC3A (80 mg/kg) or Milli-Q water alone from 3 weeks until 10 weeks of age followed by determination of **B** RBC, **C-F** Mn levels and **G-J** Zn and Cu levels in blood, brain, liver and bone (n ≥ 5 per condition). Data plotted as mean ± SD. One-Way ANOVA with Tukey’s multiple comparisons test. \*\**p* < 0.01. \**p* < 0.05

Despite its poor oral bioavailability, we hypothesized that long-term oral administration of H_3_PyC3A may reduce Mn availability from dietary sources and thus prevent Mn accumulation. *Slc30a10^-/-^* mice were allocated at weaning to receive either Milli-Q water or Milli-Q water supplemented with 80 mg/kg HCl-PyC3A in their drinking bottle for 7 weeks. Mice were sacrificed at 10 weeks and tissues and blood collected for ICP-MS analysis and determination of red blood cell counts. Mice receiving HCl-PyC3A in their drinking water failed to normalize their RBCs (**Figure 4B**), however, Mn levels in the liver, brain and blood of HCl-PyC3A-treated animals were significantly decreased compared to their age-matched untreated littermates suggesting some effect of oral chelator treatment on dietary Mn availability (**Figure 4C-F**). Nevertheless, Mn levels remained well above physiological concentrations observed in WT animals and did not fully normalize, which is the likely reason for the lack of resolution of polycythemia. Importantly, levels of other metal ions, including Zn and Cu, remained unchanged consistent with selective Mn chelation *in vivo* (**Figure 4G-J**).

We also investigated the therapeutic potential of intraperitoneal (ip) injections in the hope of increased bioavailability. In brief, *Slc30a10^-/-^* mice received 20 mg/kg of H_3_PyC3A twice daily for 2 weeks, with the treatment commencing at 3 weeks of age, and age-matched controls receiving ip injections of PBS (**Figure 5A**). We expected that efficient chelation would prevent Mn accumulation at 5 weeks. However, ip injection of H_3_PyC3A did not reduce Mn concentrations in any tissue examined (**Figure 5B-E**). This observation may support the hypothesis that oral administered H_3_PyC3A chelates Mn in the diet rather than exerting a systemic effect.

**Figure 5.**
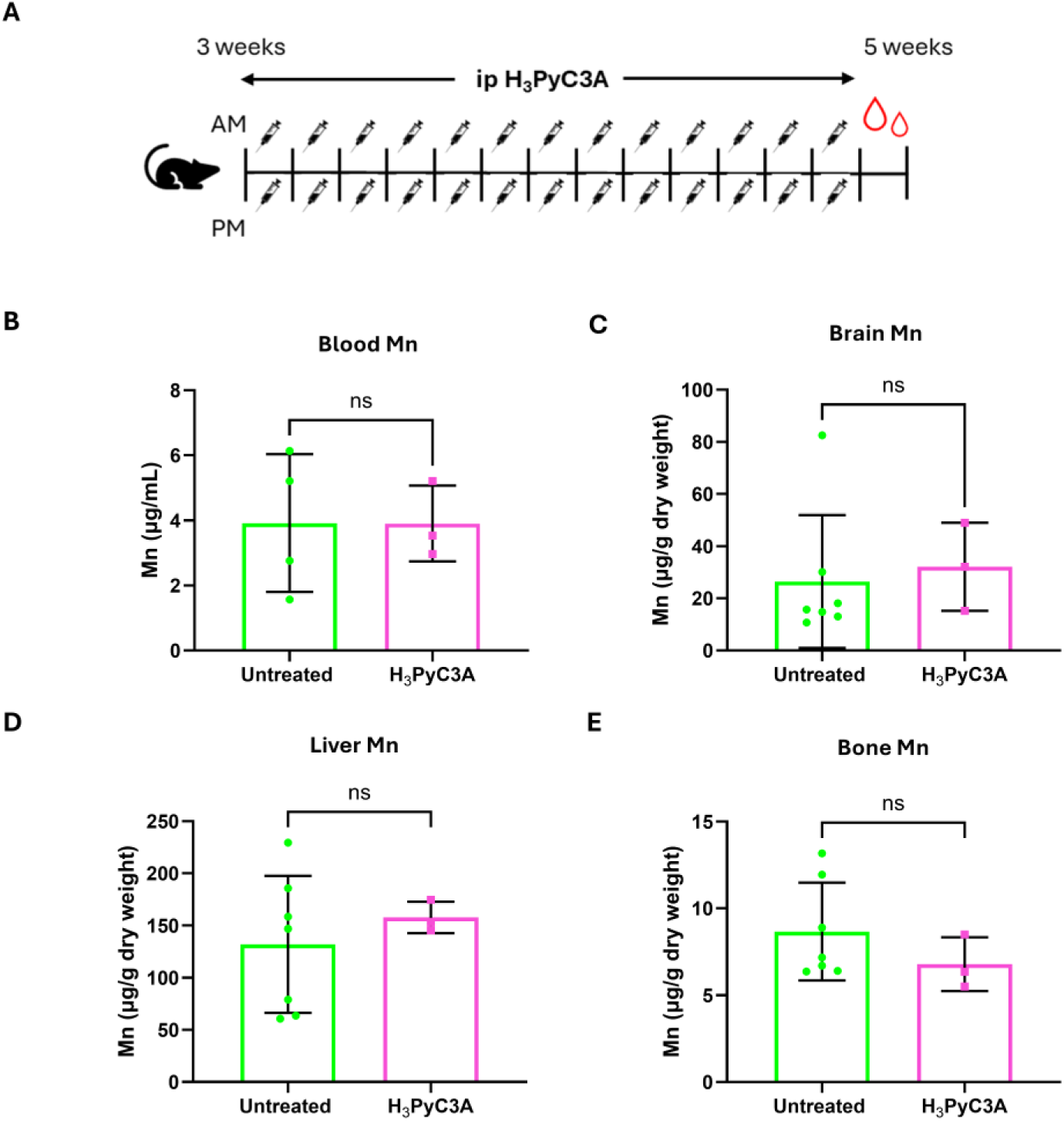
Intraperitoneal administration of HCl-PyC3A fails to reduce Mn accumulation. **A** 3-week-old Slc30a10^-/-^ mice received ip injections of either H_3_PyC3A or PBS twice a day for 2 weeks (n ≥ 5 per condition). **B-E** Mn tissue levels Data plotted as mean ± SD. One-Way ANOVA with Tukey’s multiple comparisons test. ns, not significant.

## Discussion

This study established an integrated *in vitro* to *in vivo* pipeline for the validation of novel Mn chelators, repurposed from recently developed Mn MRI contrast agents, for the treatment of Mn overload conditions (34, 48). The repurposing strategy aimed to reduce preclinical testing requirements and accelerate potential clinical translation (49). We first evaluated the Mn affinity and thermodynamic stability of complexes formed by two known Mn ligands, H_3_PyC3A and H_4_EDTA-BTA, *in vitro*. High-throughput screening of these ligands and their methyl ester derivatives using a *slc39a14^-/-^* zebrafish locomotor assay identified H_3_PyC3A as a lead compound, which was subsequently further characterized in Slc30a10 KO mice.

We were able to show that H_3_PyC3A has improved Mn binding affinity over H_4_EDTA-BTA *in vitro*. This was substantiated *in vivo* since H_3_PyC3A prominently improves the locomotor phenotype of *slc39a14^-/-^*zebrafish with almost complete normalization of whole larval Mn levels.

Despite the pronounced chelating efficacy observed in zebrafish, the high hydrophilicity of H_3_PyC3A likely contributes to its poor oral bioavailability in mice. The strong effect in zebrafish may result either from chelation of Mn in the surrounding water or from enhanced systemic uptake during early larval stages, when biological barriers are not fully developed (50). In addition to intestinal absorption, drug uptake via the skin may further increase bioavailability in larvae (51). Whilst *slc39a14^-/-^* zebrafish larvae therefore provide an ideal model for high-throughput screening of novel Mn chelators using simple locomotor assays to identify lead compounds, further studies in mammalian models are essential to characterize pharmacokinetic behavior and therapeutic potential.

The use of the methyl ester Me_3_PyC_3_A as a strategy to enhance oral bioavailability did not appear clinically advantageous (52). Pharmacokinetic studies demonstrated incomplete demethylation of Me_3_PyC_3_A, resulting in the accumulation of mono- and di-ester intermediates. These partially metabolised species would necessitate extensive preclinical safety and toxicity evaluation before further development. For this reason, Me_3_PyC_3_A was not pursued in subsequent analyses.

Interestingly, despite its poor oral bioavailability, H_3_PyC3A was able to reduce tissue Mn levels when provided continuously in the drinking water of Slc30a10 KO mice. Although red blood cell counts did not normalize, these findings indicate that either sustained chelation by maintaining a low, steady systemic exposure or reducing dietary Mn absorption through chelation of Mn in the gut may offer therapeutic benefit. The latter would suggest that chelators with limited oral absorption could serve as useful adjunctive treatments by reducing the dietary Mn load, thereby lowering systemic Mn accumulation.

Our data further demonstrate that H_3_PyC3A preferentially chelates Mn in Slc30a10 KO mice where Mn is present in supraphysiological concentrations, likely leading to excess free Mn. The *in vivo* selectivity for Mn over Zn, despite H_3_PyC3A’s higher *in vitro* stability constant for Zn, is likely to occur due to the abundance of Mn(II) in our *Slc39a14^-/-^* and *Slc30a10^-/-^* models as well as the high thermodynamic stability of the formed Mn-complexes. This suggests a form of functional selectivity under pathological conditions, where elevated Mn availability drives preferential chelation despite thermodynamic preferences observed *in vitro*. This is supported by both ICP-MS analysis showing stable Zn and Cu levels during long-term treatment with H_3_PyC3A as well as LC-MS/MS analysis of metal-PyC3A complexes demonstrating an increase in the Mn-PyC3A:Zn-PyC3A ratio in Slc30a10 KO mice that develop pronounced Mn overload. This is in keeping with observations in HMNDYT1 and HMNDYT2 patients who are effectively treated with the unselective chelator CaNa_2_EDTA (10, 14, 23, 24), which has a higher stability constant for Zn than Mn (LogK 16.5 versus 13.7). While long-term use of CaNa_2_EDTA in patients with inherited Mn transporter defects can result in low plasma Zn, regular monitoring of Zn and, in case of low blood Zn levels, Zn supplementation prevents treatment-associated deficiencies (10, 53, 54). Although an ideal Mn chelator for the treatment of Mn overload would theoretically demonstrate preferential selectivity for Mn over other physiologically relevant metals, absolute selectivity may therefore not be an essential requirement for successful therapeutic development since removal of other metals can be addressed by the use of supplements (8).

A major barrier, however, to progress Mn chelator development is the high hydrophilicity of currently used chelating agents, including H_3_PyC3A and CaNa_2_EDTA, which significantly limits their oral bioavailability. Consequently, alternative delivery strategies should be considered during drug development. Routes such as subcutaneous depot formulations, intranasal administration, or transdermal delivery may offer practical solutions to overcome short plasma half-life and poor gastrointestinal absorption (55–57). However, our findings in Slc30a10 KO mice, in which a continuous low-dose supply of H_3_PyC3A via drinking water reduced Mn burden, suggest that maintaining a stable, low systemic concentration may be sufficient to achieve meaningful chelation *in vivo*, whether this effect is caused by systemic uptake or reducing the Mn load through the diet remains to be determined.

In conclusion, our results demonstrate that H_3_PyC3A is capable of chelating Mn *in vivo*, although further optimization is required to enhance its pharmacokinetic properties and achieve clinically meaningful efficacy. The deconstruction/repurposing strategy presented here highlights a promising route for ligand design and reinvestigation of established chelators, opening new directions in coordination, bioinorganic and translational medicinal chemistry.

Moreover, the integrated *in vitro* to *in vivo* pipeline we have established, combining *slc39a14^-/-^*zebrafish for high-throughput phenotypic screening with a mammalian disease model mimicking the patient phenotype – Slc30a10 KO – mice for pharmacokinetic and proof-of-concept validation, provides a robust workflow for Mn chelator development. Given the urgent unmet clinical need for effective, ideally oral therapies to improve quality of life in individuals affected by HMNDYT1, HMNDYT2, and environmentally acquired Mn neurotoxicity, this pipeline represents a valuable foundation for future therapeutic discovery. The presented dual animal model approach has broader applicability, extending beyond Mn transport disorders to encompass other neurometabolic conditions characterized by metal dyshomeostasis.

## Author Contributions

The manuscript was written by CP, GEK, JS and KT and revised by all authors. All authors have approved the final version of the manuscript. KT and JS devised the project with critical input from RP, FMP, FCMZ, WEH, KM, PBM, AAR, JR, SWW and GEK. GEK performed crystallographic measurements and oversaw the inorganic chemistry aspects. JD synthesised the compounds, crystallised the complexes and performed data analysis with analytical and synthetic chemistry support from AM. NB and CK performed the solution studies. CD, RG-M, performed the ICP-MS analysis. SA, LEV and KT performed the zebrafish experiments. CP, HV, GF and AG performed the mouse studies. ID and NV performed the LC-MS/MS chelator assay with critical input from WEH and KM. YK performed the creatinine assay. KT, JS, LEV, and GEK performed data analysis.

## Acknowledgments

We are grateful to Prof Phil Blower at King’s College London for many fruitful discussions. We thank the EPSRC UK National Crystallography Service at the University of Southampton. We are grateful to Prof Tom Bartnikas at Brown University for sharing the Slc30a10 KO mouse model.

## References

1. Kim H, et al. Exposing the role of metals in neurological disorders: a focus on manganese. Trends Mol Med. Jul 2022;28(7):555–568. doi:10.1016/j.molmed.2022.04.011

2. Wang Y, et al. Manganese in health and disease. Nutr Res Rev. Dec 2025;38(2):900–910. doi:10.1017/S0954422425100139

3. Erikson KM, Aschner M. Manganese: Its Role in Disease and Health. Met Ions Life Sci. Jan 14 2019;19doi:10.1515/9783110527872-016

4. Baj J, et al. Consequences of Disturbing Manganese Homeostasis. Int J Mol Sci. Oct 6 2023;24(19)doi:10.3390/ijms241914959

5. Gurol KC, et al. Role of excretion in manganese homeostasis and neurotoxicity: a historical perspective. Am J Physiol Gastrointest Liver Physiol. Jan 1 2022;322(1):G79–G92. doi:10.1152/ajpgi.00299.2021

6. Tuschl K, et al. Manganese and the brain. Int Rev Neurobiol. 2013 2013;110:277–312. Not in File. doi:B978-0-12-410502-7.00013-2 [pii];10.1016/B978-0-12-410502-7.00013-2 [doi]

7. Gospe SM. Manganese Neurotoxicity and Familial Disorders of Manganese Transport. Annals of the Child Neurology Society. 2025;3(3):135–144. doi:10.1002/cns3.70015

8. Vogt H, et al. Removal of Toxic Metabolites-Chelation: Manganese Disorders. J Inherit Metab Dis. Nov 2025;48(6):e70107. doi:10.1002/jimd.70107

9. Lotz A, et al. Association of exposure to manganese and fine motor skills in welders - Results from the WELDOX II study. Neurotoxicology. Jan 2021;82:137–145. doi:10.1016/j.neuro.2020.12.003

10. Fang S, et al. Consensus of Expert Opinion for the Diagnosis and Management of Hypermanganesaemia With Dystonia 1 and 2. J Inherit Metab Dis. May 2025;48(3):e70031. doi:10.1002/jimd.70031

11. Garg D, et al. Clinical Profile and Treatment Outcomes of Hypermanganesemia with Dystonia 1 and 2 among 27 Indian Children. Mov Disord Clin Pract. Oct 2022;9(7):886–899. doi:10.1002/mdc3.13516

12. Quadri M, et al. Mutations in SLC30A10 cause parkinsonism and dystonia with hypermanganesemia, polycythemia, and chronic liver disease. Am J Hum Genet. 3/9/2012 2012;90(3):467–477. Not in File. doi:S0002-9297(12)00051-1 [pii];10.1016/j.ajhg.2012.01.017 [doi]

13. Tuschl K, et al. Syndrome of hepatic cirrhosis, dystonia, polycythemia, and hypermanganesemia caused by mutations in SLC30A10, a manganese transporter in man. Am J Hum Genet. Mar 9 2012;90(3):457–66. doi:10.1016/j.ajhg.2012.01.018

14. Tuschl K, et al. Mutations in SLC39A14 disrupt manganese homeostasis and cause childhood-onset parkinsonism-dystonia. Nat Commun. 2016;7:11601. doi:10.1038/ncomms11601

15. Chib S, Singh S. Manganese and related neurotoxic pathways: A potential therapeutic target in neurodegenerative diseases. Neurotoxicol Teratol. Sep 30 2022;94:107124. doi:10.1016/j.ntt.2022.107124

16. Hung YH, et al. Altered transition metal homeostasis in Niemann-Pick disease, type C1. Metallomics. Mar 2014;6(3):542–53. doi:10.1039/c3mt00308f

17. Martins AC, Jr., et al. New Insights on the Role of Manganese in Alzheimer’s Disease and Parkinson’s Disease. Int J Environ Res Public Health. Sep 22 2019;16(19)doi:10.3390/ijerph16193546

18. Tan J, et al. Regulation of intracellular manganese homeostasis by Kufor-Rakeb syndrome-associated ATP13A2 protein. J Biol Chem. 8/26/2011 2011;286(34):29654–29662. Not in File. doi:M111.233874 [pii];10.1074/jbc.M111.233874 [doi]

19. Zhang HT, et al. PINK1/Parkin-mediated mitophagy play a protective role in manganese induced apoptosis in SH-SY5Y cells. Toxicol In Vitro. Aug 2016;34:212–219. doi:10.1016/j.tiv.2016.04.006

20. Martins AC, et al. Manganese Induced neurodegenerative diseases and possible therapeutic approaches. Expert Review of Neurotherapeutics. 2020:null–null. doi:10.1080/14737175.2020.1807330

21. Quadri M, et al. Manganese transport disorder: Novel SLC30A10 mutations and early phenotypes. Mov Disord. 3/17/2015 2015;Not in File. doi:10.1002/mds.26202 [doi]

22. Stamelou M, et al. Dystonia with brain manganese accumulation resulting from SLC30A10 mutations: a new treatable disorder. Mov Disord. 9/1/2012 2012;27(10):1317–1322. Not in File. doi:10.1002/mds.25138 [doi]

23. Tuschl K, et al. Syndrome of hepatic cirrhosis, dystonia, polycythemia, and hypermanganesemia caused by mutations in SLC30A10, a manganese transporter in man. Am J Hum Genet. 3/9/2012 2012;90(3):457–466. Not in File. doi:S0002-9297(12)00052-3 [pii];10.1016/j.ajhg.2012.01.018 [doi]

24. Tuschl K, et al. Hepatic cirrhosis, dystonia, polycythaemia and hypermanganesaemia--a new metabolic disorder. J Inherit Metab Dis. 4/2008 2008;31(2):151–163. Not in File. doi:10.1007/s10545-008-0813-1 [doi]

25. Marti-Sanchez L, et al. Hypermanganesemia due to mutations in SLC39A14: further insights into Mn deposition in the central nervous system. Orphanet J Rare Dis. Jan 30 2018;13(1):28. doi:10.1186/s13023-018-0758-x

26. Bradberry S, Vale A. A comparison of sodium calcium edetate (edetate calcium disodium) and succimer (DMSA) in the treatment of inorganic lead poisoning. Clin Toxicol (Phila). 11/2009 2009;47(9):841–858. Not in File. doi:10.3109/15563650903321064 [doi]

27. Osterloh J, Becker CE. Pharmacokinetics of CaNa2EDTA and chelation of lead in renal failure. Clin Pharmacol Ther. Dec 1986;40(6):686–93. doi:10.1038/clpt.1986.245

28. Clement O, et al. Contrast agents in magnetic resonance imaging of the liver: present and future. Biomed Pharmacother. 1998;52(2):51–8. doi:10.1016/S0753-3322(98)80003-6

29. Drahoš B, et al. Manganese(II) Complexes as Potential Contrast Agents for MRI. European Journal of Inorganic Chemistry. 2012;2012:1975–1986. doi:10.1002/ejic.201101336

30. Miriyala S, et al. Manganese superoxide dismutase, MnSOD and its mimics. Biochim Biophys Acta. May 2012;1822(5):794–814. doi:10.1016/j.bbadis.2011.12.002

31. Wahsner J, et al. Chemistry of MRI Contrast Agents: Current Challenges and New Frontiers. Chemical Reviews. 2019/01/23 2019;119(2):957–1057. doi:10.1021/acs.chemrev.8b00363

32. Boros E, et al. MR imaging probes: design and applications. Dalton Trans. Mar 21 2015;44(11):4804–4818. doi:10.1039/c4dt02958e

33. Gupta A, et al. Applications for Transition-Metal Chemistry in Contrast-Enhanced Magnetic Resonance Imaging. Inorg Chem. May 18 2020;59(10):6648–6678. doi:10.1021/acs.inorgchem.0c00510

34. Gale EM, et al. A Manganese-based Alternative to Gadolinium: Contrast-enhanced MR Angiography, Excretion, Pharmacokinetics, and Metabolism. Radiology. Mar 2018;286(3):865–872. doi:10.1148/radiol.2017170977

35. Islam MK, et al. Manganese Complex of Ethylenediaminetetraacetic Acid (EDTA)-Benzothiazole Aniline (BTA) Conjugate as a Potential Liver-Targeting MRI Contrast Agent. J Med Chem. Apr 13 2017;60(7):2993–3001. doi:10.1021/acs.jmedchem.6b01799

36. Gale EM, et al. A Manganese Alternative to Gadolinium for MRI Contrast. Journal of the American Chemical Society. 2015/12/16 2015;137(49):15548–15557. doi:10.1021/jacs.5b10748

37. Beaumont K, et al. Design of ester prodrugs to enhance oral absorption of poorly permeable compounds: challenges to the discovery scientist. Curr Drug Metab. Dec 2003;4(6):461–85. doi:10.2174/1389200033489253

38. Banks WA. Characteristics of compounds that cross the blood-brain barrier. BMC Neurology. 2009/06/12 2009;9(1):S3. doi:10.1186/1471-2377-9-S1-S3

39. Daina A, et al. SwissADME: a free web tool to evaluate pharmacokinetics, drug-likeness and medicinal chemistry friendliness of small molecules. Sci Rep. Mar 3 2017;7:42717. doi:10.1038/srep42717

40. Yang NJ, Hinner MJ. Getting across the cell membrane: an overview for small molecules, peptides, and proteins. Methods Mol Biol. 2015;1266:29–53. doi:10.1007/978-1-4939-2272-7_3

41. Mercadante CJ, et al. Manganese transporter Slc30a10 controls physiological manganese excretion and toxicity. J Clin Invest. Dec 2 2019;129(12):5442–5461. doi:10.1172/JCI129710

42. Mills PB, et al. Genotypic and phenotypic spectrum of pyridoxine-dependent epilepsy (ALDH7A1 deficiency). Brain. Jul 2010;133(Pt 7):2148–59. doi:10.1093/brain/awq143

43. Liederer BM, Borchardt RT. Enzymes involved in the bioconversion of ester-based prodrugs. J Pharm Sci. Jun 2006;95(6):1177–95. doi:10.1002/jps.20542

44. Gascon JM, et al. Synthesis and Study of Multifunctional Cyclodextrin-Deferasirox Hybrids. ChemMedChem. Aug 20 2019;14(16):1484–1492. doi:10.1002/cmdc.201900334

45. Hassell-Hart S, et al. Synthesis and Biological Investigation of (+)-JD1, an Organometallic BET Bromodomain Inhibitor. Organometallics. 2020/01/01/ 2020;39(3):408–416. 10.1021/acs.organomet.9b00750

46. Tuschl K, et al. Loss of slc39a14 causes simultaneous manganese hypersensitivity and deficiency in zebrafish. Dis Model Mech. Jun 1 2022;15(6)doi:10.1242/dmm.044594

47. Emran F, et al. A behavioral assay to measure responsiveness of zebrafish to changes in light intensities. J Vis Exp. Oct 3 2008;(20)doi:10.3791/923

48. Poggiarelli L, et al. Manganese-Based Contrast Agents as Alternatives to Gadolinium: A Comprehensive Review. Clin Pract. Jul 25 2025;15(8)doi:10.3390/clinpract15080137

49. Zhan P, et al. Drug repurposing: An effective strategy to accelerate contemporary drug discovery. Drug Discov Today. Jul 2022;27(7):1785–1788. doi:10.1016/j.drudis.2022.05.026

50. Li Y, et al. Zebrafish as a visual and dynamic model to study the transport of nanosized drug delivery systems across the biological barriers. Colloids Surf B Biointerfaces. Aug 1 2017;156:227–235. doi:10.1016/j.colsurfb.2017.05.022

51. Lewis L, Kwong RWM. Zebrafish as a Model System for Investigating the Compensatory Regulation of Ionic Balance during Metabolic Acidosis. Int J Mol Sci. Apr 5 2018;19(4)doi:10.3390/ijms19041087

52. Rader AFB, et al. Improving oral bioavailability of cyclic peptides by N-methylation. Bioorg Med Chem. Jun 1 2018;26(10):2766–2773. doi:10.1016/j.bmc.2017.08.031

53. Tuschl K, et al. Dystonia/Parkinsonism, Hypermanganesemia, Polycythemia, and Chronic Liver Disease. 1993 1993;Not in File. doi:NBK100241 [bookaccession]

54. Tuschl K, et al. SLC39A14 Deficiency. In: Adam MP, et al, eds. GeneReviews((R)). 1993.

55. Toliyat T, et al. An extended-release formulation of desferrioxamine for subcutaneous administration. Drug Deliv. Oct 2009;16(7):416–21. doi:10.1080/10717540903141768

56. Cheng R, Kim J. Intranasal delivery of iron chelators and management of central nervous system disease. Front Pharmacol. 2025;16:1709259. doi:10.3389/fphar.2025.1709259

57. Rodrigues M, et al. Iron Chelation with Transdermal Deferoxamine Accelerates Healing of Murine Sickle Cell Ulcers. Adv Wound Care (New Rochelle). Oct 1 2018;7(10):323–332. doi:10.1089/wound.2018.0789

