## Supplemental Material for "A novel pipeline for the validation of manganese chelators for the treatment of manganese overload"

### Supplementary Methods

##### LC-MS/MS assay of PyC3A–metal complexes from dried blood spot samples LC-MS/MS

1. **Reagents and Materials**

Acetonitrile (ACN; Thermo Fisher Scientific; Cat. No. A955-4), Methanol (MeOH; Thermo Fisher Scientific; Cat. No. A456), Isopropanol (IPA; Thermo Fisher Scientific; Cat. No. A461), Milli-Q water (>18 MΩ·cm), LC Column: ACQUITY Premier BEH Amide, 1.7 µm, 2.1 × 100 mm (Waters, P/N: 186009505), 0.4mL Clear Screw PP Vial with Fixed Conical Insert (9mm Short Thread); Thermo Fisher Scientific; Cat. No. 6ESV9-04PP

##### 2. Sample Extraction Procedure

50 µL of extraction solution was added to 3 mm DBS punches in 1.5 mL Eppendorf tubes followed by gentle shaking for 30 minutes at RT. Samples were sonicated in a water bath sonicator for 15 minutes followed by centrifugation at maximum speed (~16,000 × g) for 10 minutes to pellet particulates. 40 µL of supernatant was carefully transferred into a polypropylene LC-MS vial with insert or appropriate sample plate. Note: Due to high hydrophilicity, PyC3A–metal complexes may adsorb to glass, especially borosilicate, therefore, plastic consumables were used.

##### 3. Liquid Chromatography Conditions

Chromatographic separation was carried out using an ACQUITY Premier BEH Amide column (1.7 µm, 2.1 × 100 mm; Waters) maintained at 40 °C, with the autosampler held at 4 °C. The flow rate was 0.2 mL/min and the injection volume was 3 µL.

Mobile phase A consisted of a 50:50 (v/v) mixture of water and acetonitrile containing 5 mM ammonium acetate, adjusted to pH 7.6. Mobile phase B consisted of a 95:5 (v/v) mixture of acetonitrile and water containing 5 mM ammonium acetate, adjusted to pH 7.6.

All solvents were LC–MS grade, and water was Milli-Q purified (>18 MΩ·cm). The gradient program was as follows:

TIME (MIN) A (%) B (%) DESCRIPTION

0.00	51.1 48.9 Initial conditions

| 3.00 | 51.1 | 48.9 | Isocratic hold |
| --- | --- | --- | --- |
| 7.00 | 77.8 | 22.2 | Linear gradient |
| 9.00 | 77.8 | 22.2 | Hold |
| 9.01 | 51.1 | 48.9 | Return to initial |
| 15.00 | 51.1 | 48.9 | Re-equilibration |
| Wash | solvents used | by the | autosampler consisted of a strong needle wash comprising |

acetonitrile:methanol:isopropanol:water (25:25:25:25, v/v/v/v) and a weak wash comprising acetonitrile:water (95:5, v/v).

Prior to analysis, the column was equilibrated for at least 30 column volumes (approximately 1 h) under initial conditions. A minimum of five blank or standard injections were performed before sample analysis to stabilise the HILIC stationary phase and ensure reproducible retention times.

##### 4. Mass Spectrometry Conditions

Detection was performed using a Xevo TQ-S triple quadrupole mass spectrometer (Waters) operated in negative electrospray ionisation mode.

Source parameters were as follows: capillary voltage 2.0 kV, source offset 30 V, desolvation temperature 450 °C, desolvation gas flow 800 L/hr, cone gas flow 15 L/hr. and nebuliser gas pressure 7 bar.

Quadrupole and collision cell settings were maintained at LM resolution 2.80, HM resolution 14.90, ion energy 0.5 for both stages and collision gas flow 0.15 mL/min. Compound-specific cone voltages and collision energies were used for all monitored transitions.

##### 5. Multiple Reaction Monitoring (MRM) Transitions

PyC3A–Mn²⁺:

| Precursor (m/z) | Product (m/z) | Cone (V) | CE (eV) |
| --- | --- | --- | --- |
| 431.0766 | 141.1515 | 62 | 38 |
|  | 267.0686 | 62 | 24 |
|  | 295.0103 | 62 | 28 |
| PyC3A–Co²⁺: |  |  |  |
| Precursor (m/z) | Product (m/z) | Cone (V) | CE (eV) |
| 435.0873 | 110.9446 | 68 | 62 |
|  | 286.0312 | 68 | 22 |
|  | 299.0260 | 68 | 26 |

| PyC3A–Cu²⁺: |  |  |  |
| --- | --- | --- | --- |
| Precursor (m/z) | Product (m/z) | Cone (V) | CE (eV) |
| 439.0873 | 183.0429 | 28 | 44 |
|  | 309.0769 | 28 | 26 |
|  | 351.1115 | 28 | 12 |
| PyC3A–Zn²⁺: |  |  |  |
| Precursor (m/z) | Product (m/z) | Cone (V) | CE (eV) |
| 440.0873 | 183.1165 | 40 | 38 |
|  | 276.0851 | 40 | 28 |
|  | 291.0306 | 40 | 28 |
| Free PyC3A: |  |  |  |
| Precursor (m/z) | Product (m/z) | Cone (V) | CE (eV) |
| 378.1424 | 88.0116 | 48 | 32 |
|  | 164.4629 | 48 | 24 |
|  | 245.1031 | 48 | 22 |
|  | 274.1446 | 48 | 18 |
|  | 290.1933 | 48 | 18 |

##### 6. Notes and Considerations

HILIC performance was dependent on appropriate column conditioning, and stable retention times were achieved through extended equilibration and dummy injections prior to sample analysis.

### Supplementary Figures and Tables

##### Synthesis of H3PyC3A

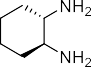

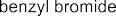

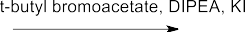

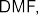

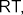

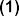

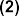
**A**

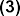

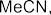

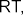

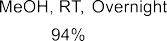

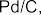

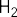

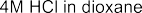

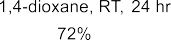

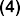

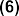
**B**

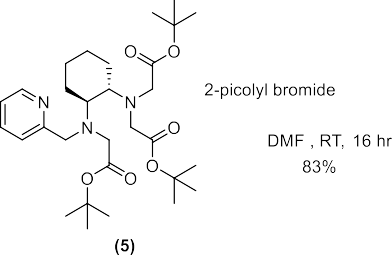

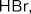

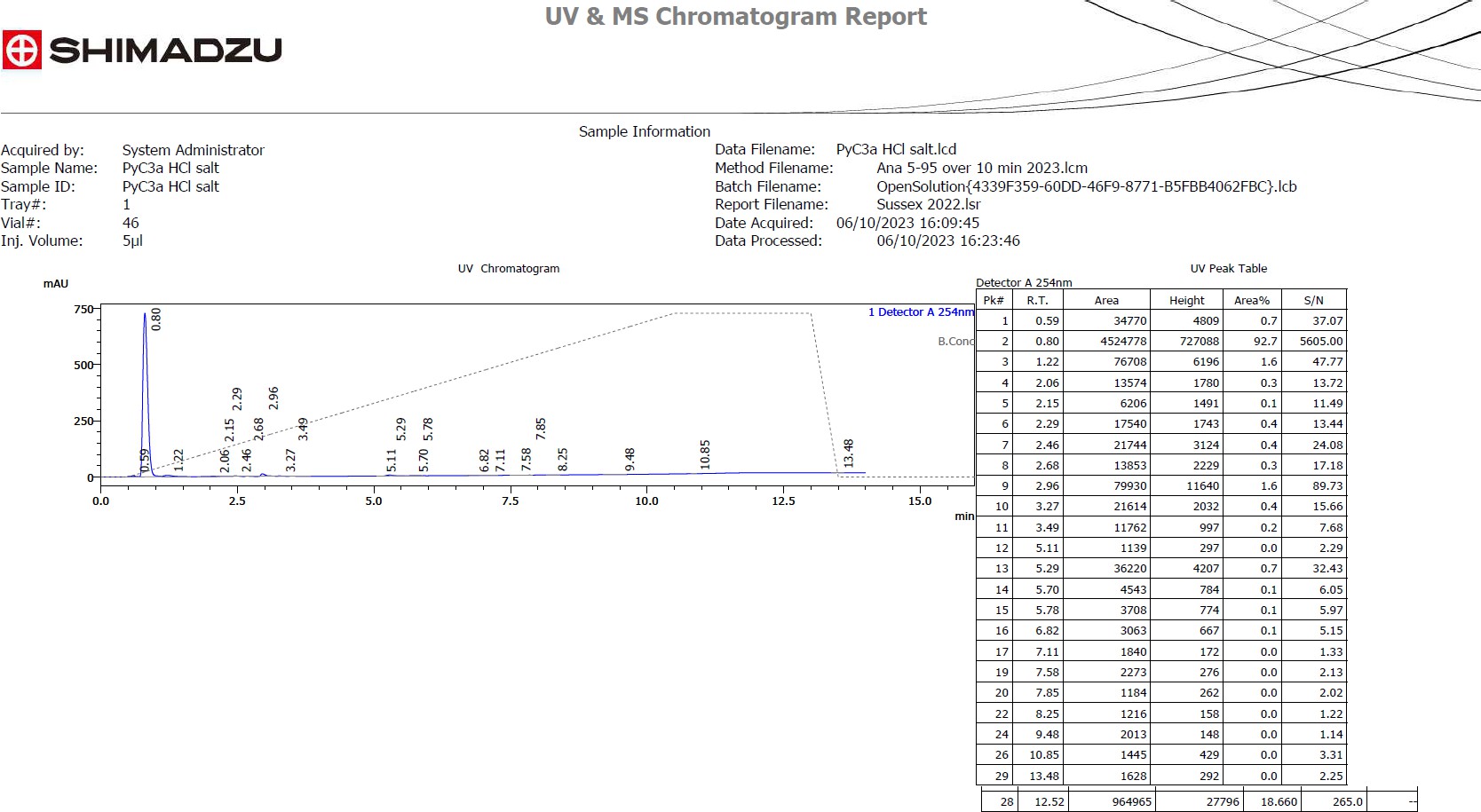

**Figure S1. A** Synthetic route for synthesis of H3PyC3A (6).

*N^1^-Benzylcyclohexane-1,2-diamine* (2):

The title compound was synthesised using the previously reported literature method.^1^ Yellow oil (1.72 g, 8.42 mmol, 96%); HRMS + pESI) calcd C13H21N2 [M + H]^+^: 205.1699, observed: 205.1693.

*di-tert-Butyl 2,2'-((2-(benzyl(2-(tert-butoxy)-2-oxoethyl)amino)cyclohexyl)azanediyl)diacetate* (3):

The title compound was synthesised using the previously reported literature method.^1^ Light yellow oil (1.17 g, 2.13 mmol, 37%); (HRMS + pESI) calcd C31H51N2O6 [M + H]^+^: 547.3742, observed: 547.3734, LC-MS: Rt = 15.97, 100%, m/z: 547.30.

*di-tert-Butyl 2,2'-((2-((2-(tert-butoxy)-2-oxoethyl)amino)cyclohexyl)azanediyl)diacetate* (4):

The title compound was synthesised using the previously reported literature method.^1^ (HRMS + pESI) calcd C24H45N2O6 [M + H]^+^: 457.3272, observed: 457.3253.

*di-tert-Butyl-2,2'-((-2-((2-(tert-butoxy)-2-oxoethyl)(pyridin-2-ylmethyl)amino)cyclohexyl)azanediyl)diacetate* (5):

The title compound was synthesised using the previously reported literature method using 2-(bromomethyl)pyridine hydrobromide in place of 2-(chloromethyl)pyridine hydrochloride. Light brown oil (860 mg, 1.57 mmol, 83%); (HRMS + pESI) calcd C30H50N3O6 [M + H]^+^: 548.3694, observed: 548.3685; LC-MS: Rt = 15.41, 86%, m/z: 548.30

*2,2'-((2-((Carboxymethyl)(pyridin-2-ylmethyl)amino)cyclohexyl)azanediyl)diacetic acid (PyC3A)* (6):

To compound 5 (860 mg, 1.57 mmol) in 1,4-dioxane (5 mL) was added 4M HCl in dioxane (5 mL). The reaction mixture was stirred at room temperature for 12 hours. The off-white precipitate was collected via filtration and washed with 1,4-dioxane (2 x 10 mL) and Et2O (2 x 10 mL) yielding the final product as light brown solid (428 mg, 72%); (HRMS + pESI) calcd C18H26N3O6 [M + H]^+^: 380.1816, observed: 380.1821; LC-MS: Rt = 0.68, 93%, m/z = 380.10

**B** Chromatogram of HCl-PyC3A

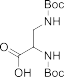

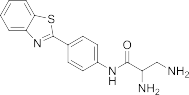

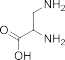

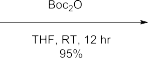

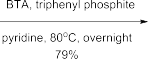

##### Synthesis of H3EDTA-BTA

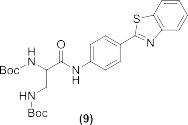

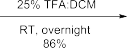

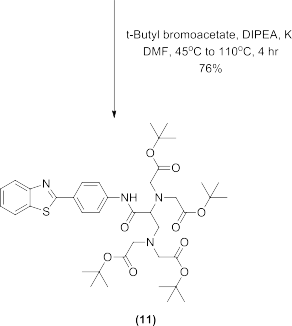

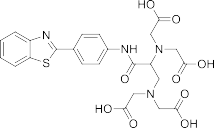

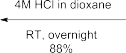
***Figure S2.*** *Synthetic route for synthesis of* ***EDTA-BTA*** *(12)*

*2,3-bis((tert-Butoxycarbonyl)amino)propanoic acid* (8):

The title compound was synthesised using the previously reported literature method.^2^ Colorless solid (3.69 g, 12.10 mmol, 95%); (HRMS + pESI) calcd C13H25N2O6 [M + H]^+^: 205.1707, observed: 305.1703.

*di-tert-Butyl (3-((4-(benzothiazol-2-yl)phenyl)amino)-3-oxopropane-1,2-diyl)dicarbamate* (9):

The title compound was synthesised using the previously reported literature method.^2^ Colorless solid (4.63 g, 9.02 mmol, 79%); (HRMS + pESI) calcd C26H33N4O5S1 [M + H]^+^: 513.2166, observed: 513.2158;

LC-MS: Rt = 23.77, 100%, m/z: 513.15.

*2,3-Diamino-N-(4-(benzo[d]thiazol-2-yl)phenyl)propanamide* (10):

The title compound was synthesised using the previously reported literature method and isolated as its TFA salt.^[3]^ Yellow solid (2.19 g, 3.34 mmol, 86%); (HRMS + pESI) calcd C16H17N4O1S1 [M + H]^+^: 313.1118, observed: 313.1104; LC-MS: Rt = 10.27 min, 96%, m/z: 312.85

*[1-(4-Benzothiazol-2-yl-phenylcarbamoyl)-2-(bis-tert-butoxycarbonylmethylamino)ethyl]-tert-butoxycarbonyl-methylamino) acetic acid tert-butyl ester* (11):

The title compound was synthesised using the previously reported literature method.^2^ Yellow oil (895 mg, 1.16 mmol, 76%); LC-MS: Rt = 4.35 min, 93%, m/z: 769.40.

*[{1-(4-Benzothiazol-2-yl-phenylcarbamoyl)-2-(biscarboxymethylamino)ethyl]carboxymethylamino}acetic acid] (BTA-EDTA)* (13):

To compound 11 (895 mg, 1.16 mmol) in 1,4-dioxane (5 mL) was added 4 M HCl in dioxane (5 mL) was added. The resulting solution was stirred at room temperature until all starting material was consumed as confirmed by LC-MS analysis; (HRMS + pESI) calcd C24H25N4O9S1 [M + H]^+^: 545.1303, observed: 545.1338; LC-MS: Rt = 2.78 min, 99%, m/z: 545.20.

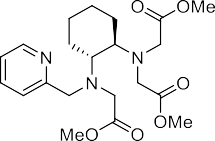

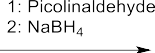

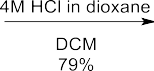

##### Synthesis of Me3PyC3A

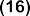

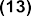

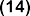

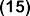

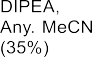
***Figure S3.*** *Synthetic route for synthesis of* ***Me3PyC3A***

*tert-Butyl-2-((pyridin-2-ylmethyl)amino)cyclohexyl)carbamate* (14):

The title compound was synthesised using a previously reported procedure.^3^ Colourless oil (291 mg, 0.95 mmol, 95%); used in the next step without further purification.

*N1-(Pyridin-2-ylmethyl)cyclohexane-1,2-diamine hydrochloride* (15):

Compound (14) (2.20 g, 7.2 mmol) was suspended in DCM (30 mL) and 4M HCl in dioxane (10 mL) was added slowly over 5 minutes. The reaction was stirred at RT overnight. The colorless precipitate was collected via gravity filtration, washed with DCM (3 x 10 mL) and dried in air to yield (15) as a colorless solid (1.79 g, 5.59 mmol, 79%); (HRMS + pESI) calcd C12H20N3 [M + H]^+^: 206.1652, observed: 206.1652.

*Dimethyl 2,2'-((-2-((2-Methoxy-2-oxoethyl)(pyridin-2-ylmethyl)amino)cyclohexyl)azanediyl)diacetate*

(16):

Compound (15) (945 mg, 3 mmol) and DIPEA (3.39 mL, 19.5 mmol, 6.5 eq.) were stirred in dry MeCN (20 mL) under a Ar atmosphere at RT. To this solution methyl bromoacetate (0.99 mL, 10.5 mmol, 3.5 eq.) was added dropwise and the reaction heated at 80 ^o^C overnight. Reaction mixture cooled to RT and diluted with D.I water (40 mL) then extracted with DCM (3 x 30 mL). The combined organics were washed with sat. Na2CO3 solution (2 x 20 mL), dried over MgSO4 and concentrated to dryness. The crude product was purified column chromatography (Hex:EtOAc + 0.5% NEt3, 1:1 to 1:4) to give (16) as a light brown solid (358 mg, 0.85 mmol, 28%); ^1^H NMR (600 MHz, dmso) δ 8.44 – 8.40 (m, 1H), 7.76 – 7.68

(m, 2H), 7.22 (d, *J* = 6.3 Hz, 1H), 4.05 (d, *J* = 14.4 Hz, 1H), 3.71 – 3.65 (m, 1H), 3.54 - 3.52 (m, 15H), 2.76

(td, *J* = 10.6, 3.3 Hz, 1H), 2.62 (d, *J* = 10.4 Hz, 1H), 1.96 – 1.85 (m, 3H), 1.60 (d, *J* = 12.4 Hz, 3H), 1.10 –

0.96 (m, 3H); ^13^C NMR (151 MHz, dmso-d6) δ 172.5, 148.5, 136.8, 123.8, 122.4, 63.0, 61.8, 56.0, 52.2,

52.0, 51.6, 28.1, 27.3, 25.6; (HRMS + pESI) calcd C21H32N3O6 [M + H]^+^: 422.2286, observed: 422.2263;

LC-MS: Rt = 12.90, 93%, m/z: 422.30.

##### Synthesis of Me4EDTA-BTA

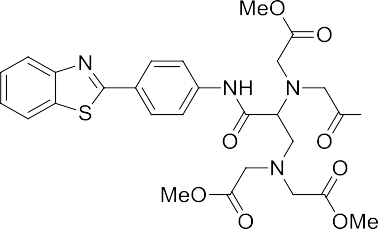

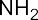

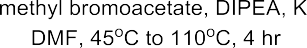

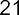

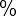

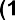

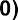

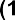

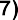

***Figure S4.*** *Synthetic route for synthesis of* ***Me4EDTA-BTA***

*[1-(4-benzothiazol-2-yl-phenylcarbamoyl)-2-(bis-tert-butoxycarbonylmethylamino)ethyl]-tert-butoxycarbonyl-methylamino) acetic acid methyl ester (17)*

Compound (17) was synthesised using the same reported literature method as compound (11) with methyl bromoacetate in place of tert-butyl bromoacetate.^[3]^ Brown oil (62 mg, 21%); (HRMS + pESI) calcd C28H33N4O9S1 [M + H]+ : 601.1963, observed: 601.1964 LC-MS: Rt = 24.12, Purity = 96%, m/z: 601.10.

**Compound characterisation**

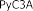

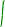

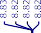

## A

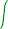

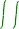

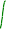

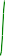

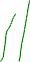

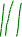

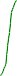

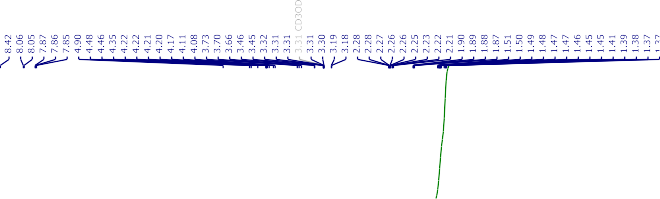

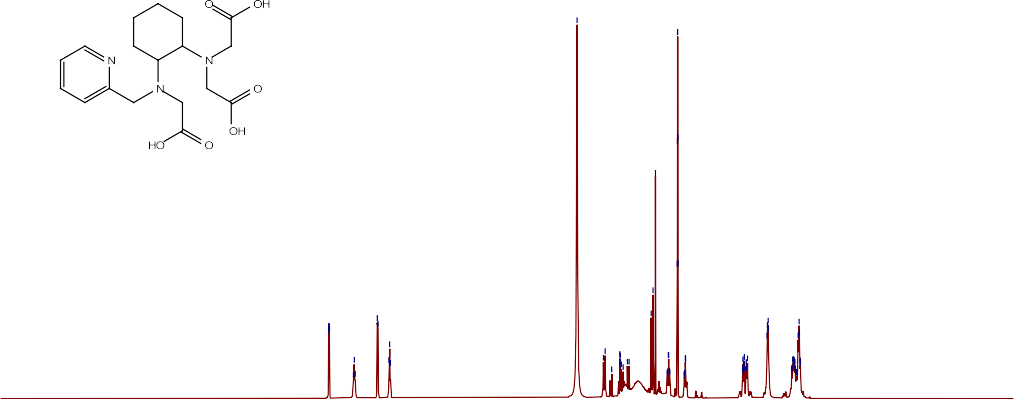

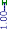

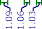

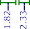

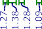

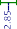

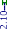

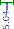

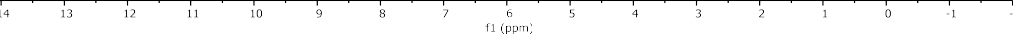

## B

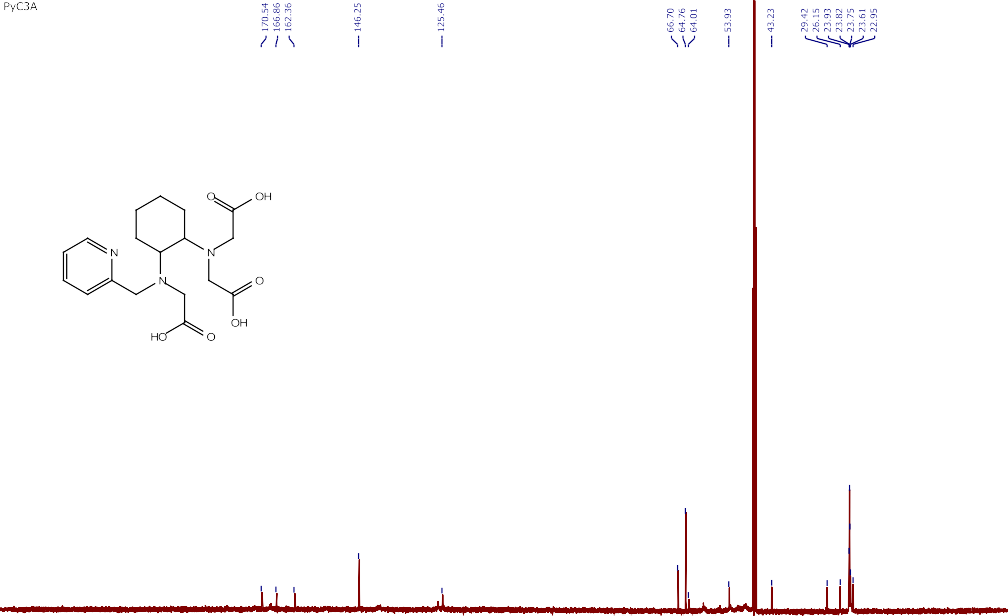

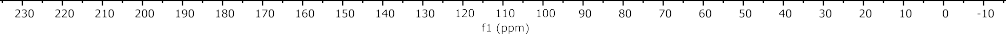

***Figure S5. A*** *^1^H-NMR and* ***B*** *^13^C-NMR spectrum of* ***PyC3A*** *(6).*

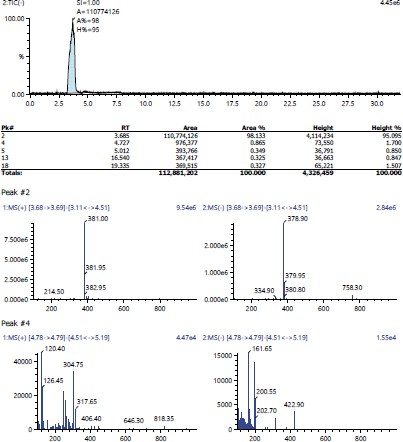

***Figure S6.*** *LC-MS trace of* ***H3PyC3A*** *(6).*

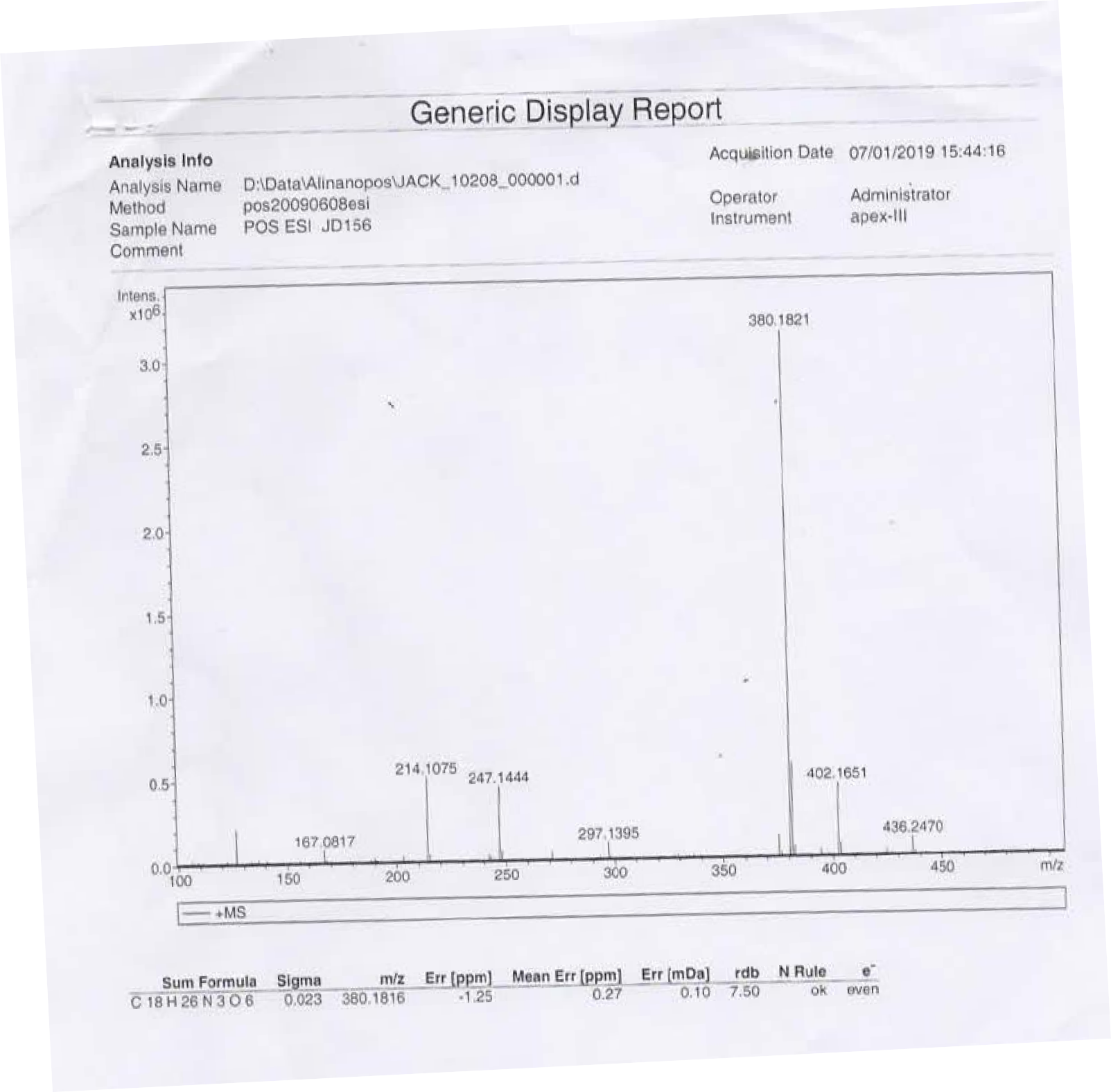

***Figure S7.*** *ESI-HRMS spectrum of* ***H3PyC3A*** *(6).*

## A

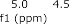

## B

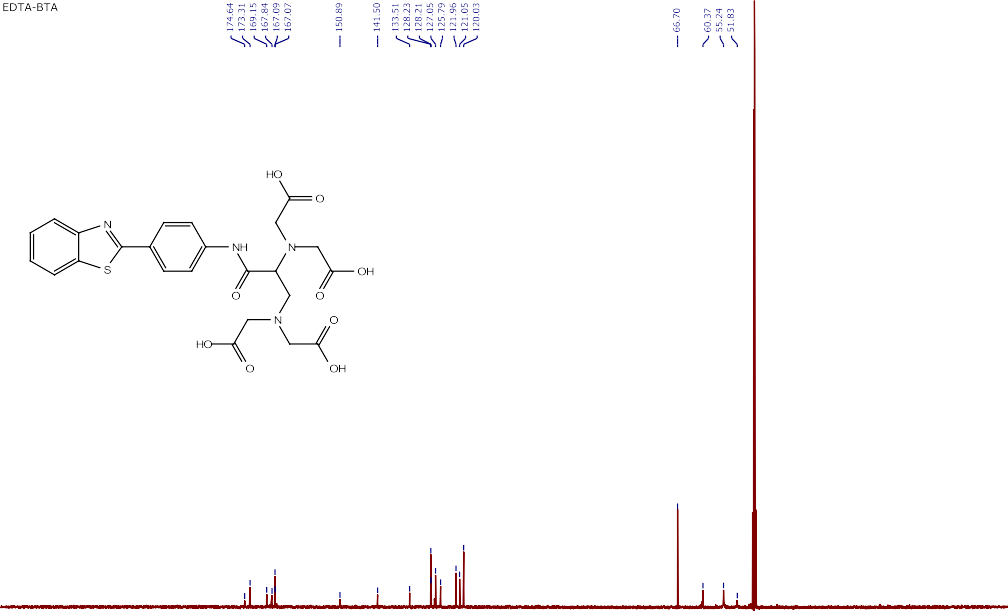

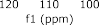

***Figure S8. A*** *^1^H-NMR* ***B*** *and ^13^C-NMR spectrum of* ***H4EDTA-BTA*** *(12).*

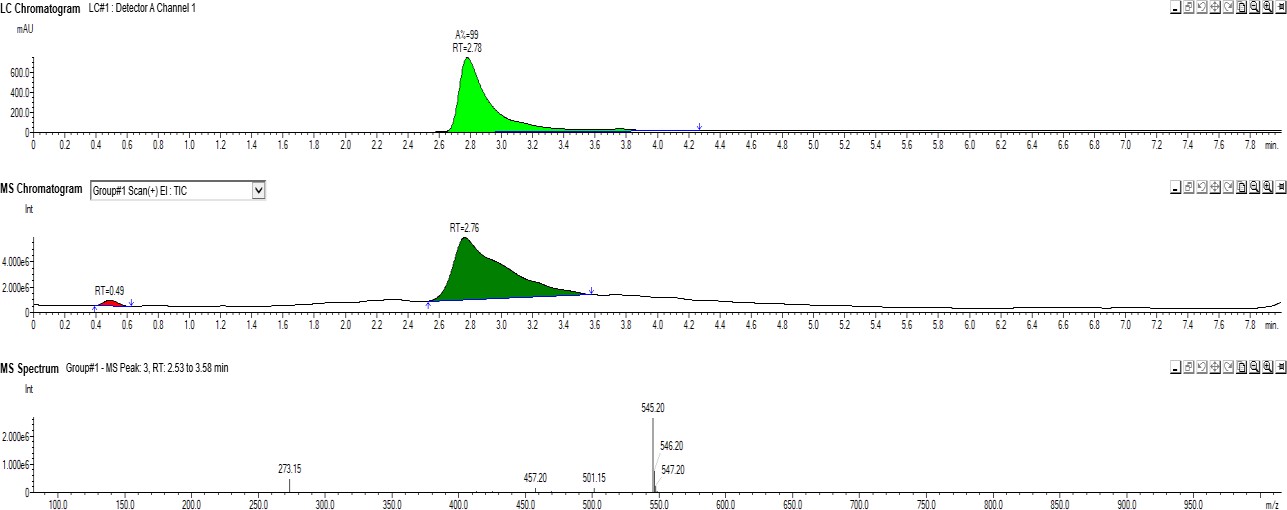

***Figure S9.*** *LC-MS trace of* ***H4EDTA-BTA*** *(12).*

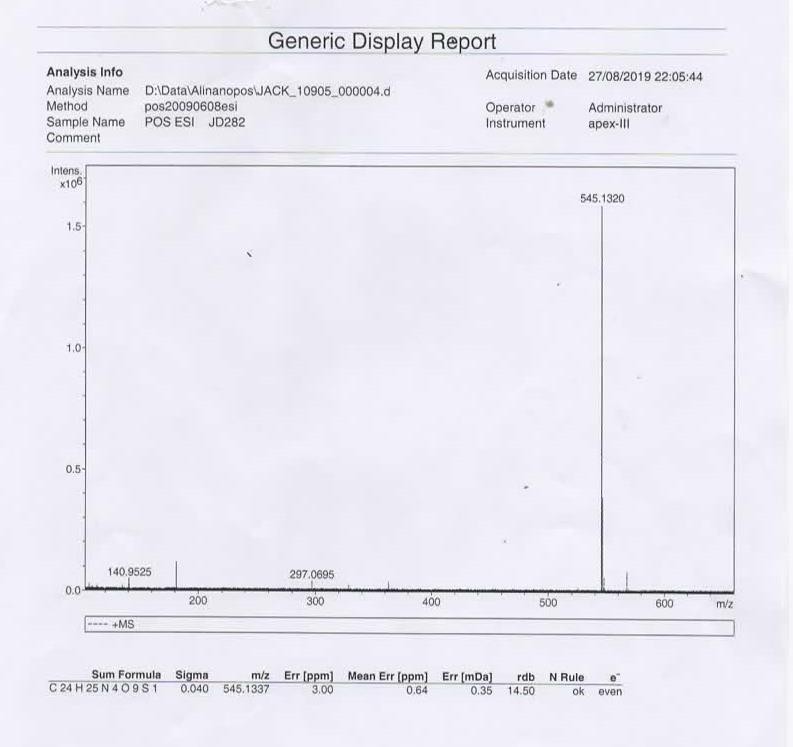

***Figure S10.*** *ESI-HRMS spectrum of* ***H4EDTA-BTA*** *(12).*

## A

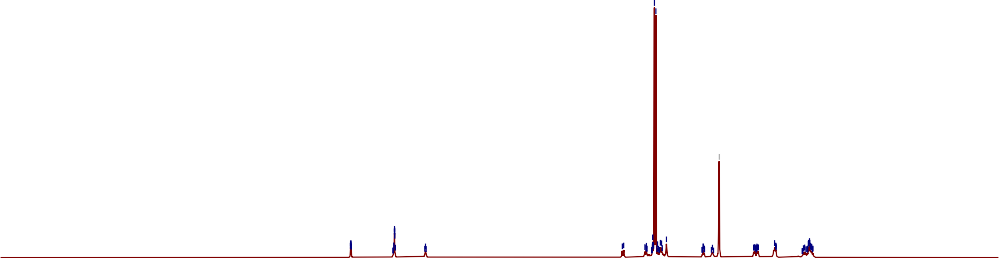

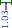

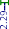

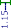

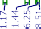

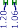

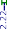

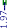

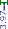

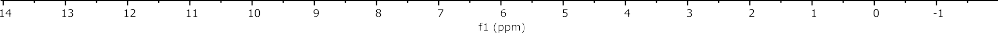

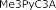

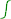

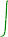

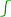

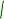

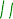

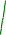

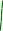

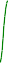

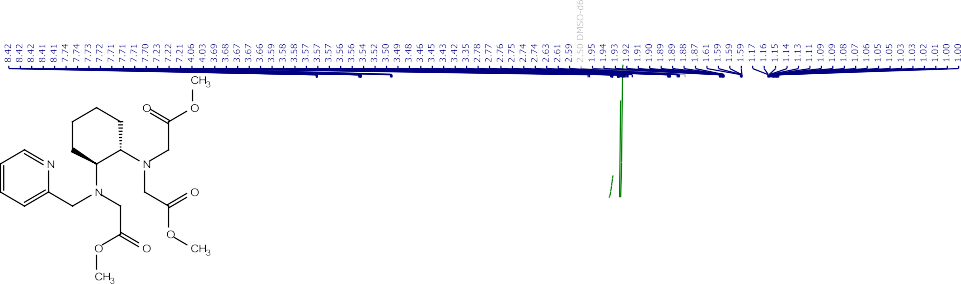

## B

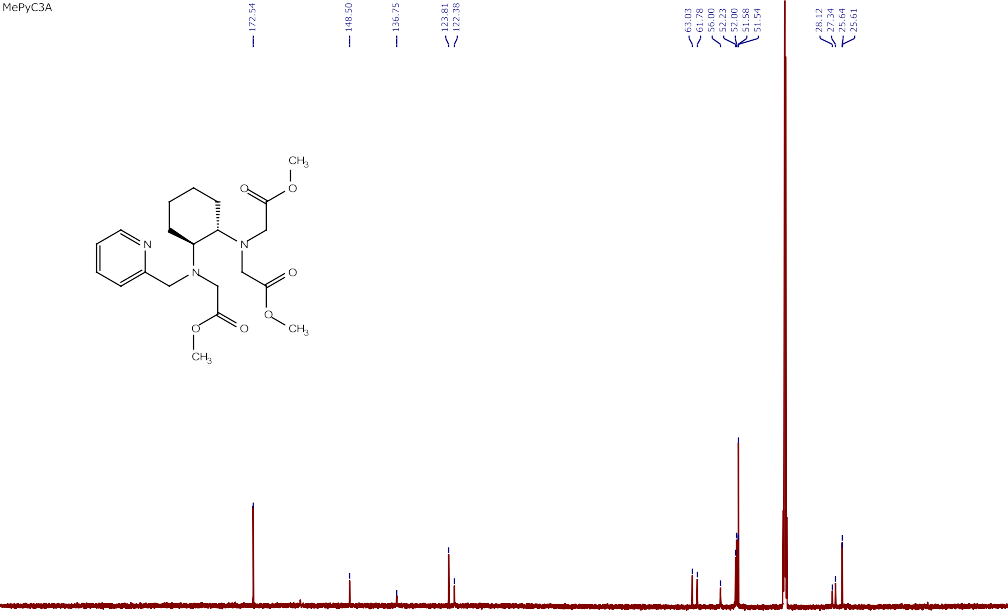

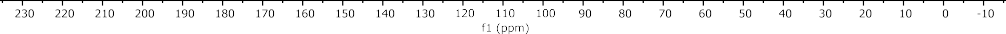

***Figure S11. A*** *^1^H-NMR and* ***B*** *^13^C-NMR spectrum of* ***Me3PyC3A*** *(16).*

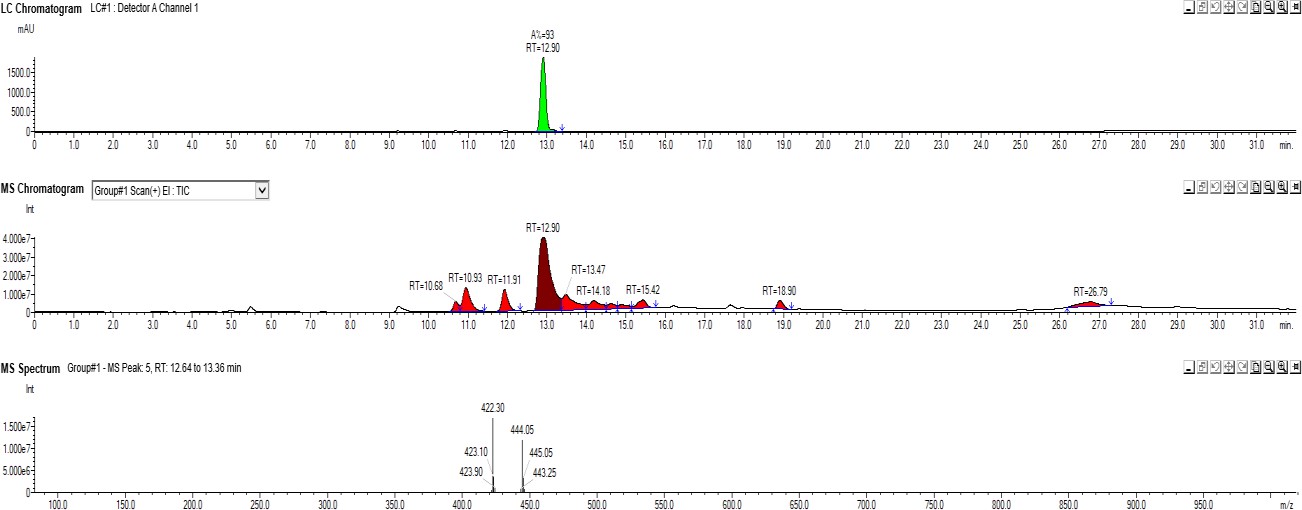

***Figure S12****. LC-MS trace of* ***Me3PyC3A*** *(16).*

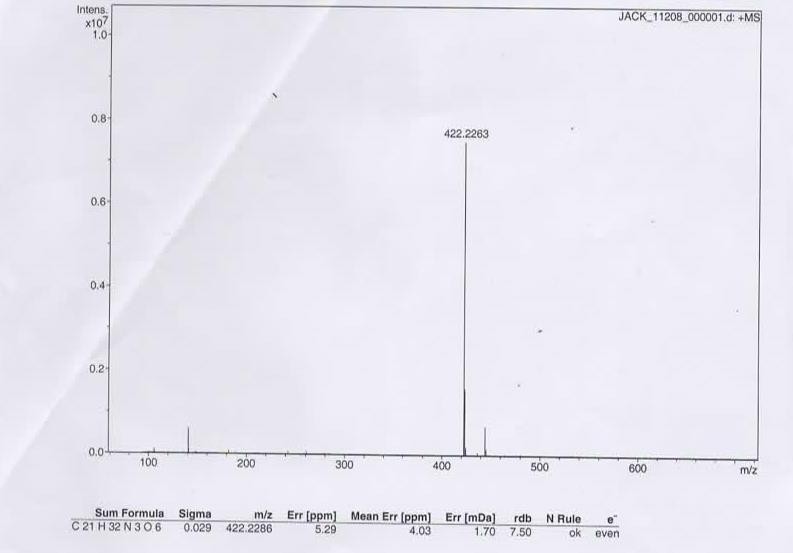

***Figure S13.*** *ESI-HRMS spectrum of* ***Me3PyC3A*** *(16).*

## A

## B

***Figure S14. A*** *^1^H-NMR* ***B*** *and ^13^C-NMR spectrum of* ***Me4EDTA-BTA*** *(17).*

***Figure S15.*** *LC-MS trace of* ***Me4EDTA-BTA*** *(17).*

***Figure S16.*** *ESI-HRMS spectrum of* ***Me4EDTA-BTA*** *(17).*

##### Crystallographic analysis

Crystallographic data were obtained by collaborators at EPSRC UK National Crystallography Service at the University of Southampton as previously described.^4^

*Complexation study of Me3PyC3A*

Me3PyC3A (50 mg. 0.12 mmol) and MnCl2.4H2O (24 mg, 0.12 mmol) were suspended in MeOH (5 mL) and stirred at room temperature for 24 h. The solution was analysed by HRMS and left to evaporate to dryness. Slow evaporation of the oily residue from dichloromethane led to isolation of colorless crystals.

We attempted to gain more evidence for this system's solid and solution behavior by monitoring an equimolar reaction between Me3PyC3A and MnCl2.4H2O in methanolic solutions. Electrospray ionisation mass spectrometry (ESI-MS) revealed peaks corresponding to [Mn(Me3PyC3A)Cl^+^] and the free ligand. Slow evaporation led to the isolation of tiny, colorless crystals. ESI-MS of the resulting powder showed the presence of the main peak corresponding to Mn(Me2PyC3A)^+^, indicating the loss of one methyl ester group to afford the corresponding monoacid. Crystallographic characterisation of this solid identified a tetrameric [Mn4(Me2PyC3A)3Cl3][MnCl4] star-like species, supporting the ESI-MS findings. The weakly diffracted crystals, even in a Synchrotron source, are only presented herein to support the evidence observed in ESI-MS. From these data, it is evident that three metal ligands [Mn(Me2PyC3A)] surround a Mn^2+^ ion bridged via a chloride and oxygen of the carboxylic acid groups from the hydrolyzed ligand (**Figure S17**).

***Figure S17.*** *ESI-HRMS spectrum of* ***A Me3PyC3A*** *ligand, showing main peak corresponding to [Me3PyC3A+H]^+^ = 422.2263,* ***B Me3PyC3A*** *+ MnCl2 in MeOH solution showing peaks corresponding to;*

*[Me3PyC3A+Na]^+^ = 444.2101 and [Mn(Me3PyC3A)Cl]^+^ = 511.1253,* ***C*** *Colorless crystals showing main peak corresponding to [[Mn(Me2PyC3A)]+H]^+^ = 461.1353.* ***D*** *Crystallographic structure of [Mn4(Me2PyC3A)3Cl3]^2+^. The tetrameric [Mn4(Me2PyC3A)3Cl3]^2+^ species formed in solid state with MnCl42-counter ions and lattice solvent molecules omitted for clarity. Colour code: Mn (purple); O (red), N (light blue), C (grey), Cl (light green).*

*Mn-EDTA-BTA polymetallic species*

EDTA-BTA (100 mg, 0.19 mmol) and MnCl2.4H2O (37 mg, 0.19 mmol) were suspended in MeOH (5 mL) and the pH was adjusted to 6 with 1M NaOH solution. The mixture was stirred at room temperature overnight and the formed precipitate was collected via filtration. Crystalline material was obtained by the slow diffusion of acetone into a concentrated aqueous solution.

X-ray quality crystals of purported Mn-EDTA-BTA were obtained by suspending H4EDTA-BTA and MnCl2.4H2O in MeOH and adjusting the pH to 6.5 with 1M NaOH. The resulting precipitate was collected, and vapor diffusion (acetone-H2O) crystallization afforded colourless needles after four weeks. The crystallographic data revealed co-crystallisation of a one dimensional polymeric chain and a discrete tetrameric species. Negative ESI mass spectrometric analysis of the crystals showed that the expected [Mn(EDTA-BTA)(H2O)^2-^] entity remains the main species in solution (**Figure S18**).

***Figure S18.*** *Crystallographic structure of Mn(EDTA-BTA)(H2O)^2-^****A*** *Projection of the 1D polymeric chain consisting of* ***Mn(EDTA-BTA)(H2O)^2-^*** *bridged by Mn(H2O)42+ units and* ***B*** *The co-crystallised tetrameric entity formed from two* ***[Mn(EDTA-BTA)]^2-^*** *units and two [Mn(H2O)4]^2+^ units (lower) Color code: Mn (purple); O (red), N (light blue), C (gray), Cl (light green)* ***C*** *Negative ESI-HRMS spectrum of crystals showing main peak corresponding to [Mn(EDTA-BTA)]^2-^ = 596.0404 species.*

***Table S1****. Crystal data and structure refinement for Mn-EDTA-BTA polymetallic species and [Mn4(Me2PyC3A)3Cl3][MnCl4].*

| Empirical formula | C27H40Mn2N4O17S | C1296H1512N72O288Cl144Mn108 |
| --- | --- | --- |
| Formula weight | 834.57 | 33744.01 |
| Temperature/K | 100(2) | 100.15 |
| Crystal system | triclinic | trigonal |
| Space group | P-1 | R-3 |
| a/Å | 7.5812(5) | 17.9897(8) |
| b/Å | 9.3943(6) | 17.9897(8) |
| c/Å | 24.9824(17) | 94.240(4) |
| α/° | 86.277(5) | 90 |
| β/° | 88.868(5) | 90 |
| γ/° | 81.387(5) | 120 |
| Volume/Å^3^ | 1755.4(2) | 26413(3) |
| Z | 2 | 1 |
| ρcalcg/cm^3^ | 1.579 | 2.121 |
| μ/mm^-1^ | 0.859 | 1.574 |
| F(000) | 864.0 | 17244.0 |
| Crystal size/mm^3^ | 0.13 × 0.03 × 0.01 | 0.1 × 0.08 × 0.01 |
| Radiation | MoKα (λ = 0.71075) | ? (λ = 0.6889) |
| 2Θ range for data collection/° | 4.394 to 46.514 | 3.038 to 45 |
| Index ranges | -8 ≤ h ≤ 8, -10 ≤ k ≤ 10, -27 ≤ l ≤ 27 | 19 ≤ h ≤ 19, -19 ≤ k ≤ 19, -104 ≤ l ≤ 104 |
| Reflections collected | 27457 | 93667 |
| Independent reflections | 5057 [Rint = 0.1117, Rsigma = 0.0735] | 8430 [Rint = 0.1983, Rsigma = 0.0900] |
| Data/restraints/parameters | 5057/154/465 | 8430/0/671 |
| Goodness-of-fit on F^2^ | 1.038 | 2.247 |
| Final R indexes [I>=2σ (I)] | R1 = 0.1337, wR2 = 0.3163 | R1 = 0.2261, wR2 = 0.5595 |
| Final R indexes [all data] | R1 = 0.1601, wR2 = 0.3395 | R1 = 0.2649, wR2 = 0.5780 |
| Largest diff. peak/hole / e Å^-3^ | 7.32/-3.56 | 5.68/-1.45 |

***Figure S19:*** *Titration curves of* ***H3PyC3A*** *and the metal ion containing systems* ***deprotonation and stability constants***

***Table S2:*** *Deprotonation constants of* ***H3PyC3A****.*

| pK1 | 2.62(9) |
| --- | --- |
| pK2 | 2.83(8) |
| pK3 | 4.02(6) |
| pK4 | 6.59(5) |
| pK5 | 11.59(5) |

***Table S3:*** *Stability constants of the Mn(II) and Zn(II)* ***H3PyC3A*** *complexes.*

|  | Mn(II) | Zn(II) |
| --- | --- | --- |
| [MLH] | 19.57(2) | 21.88(3) |
| [ML]^−^ | 15.54(4) | 18.45(4) |

***Figure S20****: Titration curves of* ***H4EDTA-BTA*** *and the metal ion containing systems in DMSO:water mixture.*

***Table S4:*** *Deprotonation constants of* ***H3PyC3A*** *and* ***H4EDTA-BTA*** *when studied in a DMSO:water system.*

|  | **H3PyC3A** | **H4EDTA-BTA** |
| --- | --- | --- |
| **pK1** | 4.19(9) | 4.59(9) |
| **pK2** | 4.81(6) | 5.43(8) |
| **pK3** | 6.35(6) | 9.87(6) |
| **pK4** | 11.19(4) | - |

***Table S5:*** *Stability constants of the metal complexes studied in DMSO:water mixture.*

|  | **H3PyC3A** | **H4EDTA-BTA** | |
| --- | --- | --- | --- |
|  | **Mn** | **Mn** | **Zn** |
| **[MLH2]^+/0^** | 23.80(2) | 21.86(13) | 22.30(3) |
| **[MLH]^0/^**^−^ | 19.58(3) | 17.60(6) | 17.72(6) |
| **[ML]**−**/2**− | 13.87(5) | 9.85(20) | 8.07(13) |

***Figure S21****:* ***A-B*** *Species distribution in H2O of H3PyC3A with* ***A*** *Mn(II) and* ***B*** *Zn(II).* ***C*** *Species distribution in DMSO:H2O of H3PyC3A (70:30) with Mn(II).* ***D-E*** *Species distribution of H4EDTA-BTA in DMSO:H2O (70:30) with* ***D*** *Mn(II) and* ***E*** *Zn(II).*

**E D T A - B T A**

**6** U n e x p o s e d

M n C l_2_ 5 0 μ M

M n C l_2_ 5 0 μ M + E D T A - B T A

**s e c o n d s /m in u te**

**4**

**2**

**0**

**4	5	6	7**

**d a y s p o s t fe r t iliz a t io n**

***Figure S22.*** *Locomotor activity is unchanged upon* ***H4EDTA-BTA*** *treatment in MnCl2 exposed slc39a14^-/-^zebrafish. MnCl2 (50 µM) and H4EDTA-BTA (10 µM) were added from 2 dpf. Average day activity (mean±SEM) is plotted from 4 to 7 dpf (n=24 per condition).*

***Figure S23.*** *Levels of Zn (C), Fe (D), and Ca (E) are not significantly reduced by cardiac injection of H3PyC3A (n≥5 per group). Student’s t-test.*P<0.05, ns=not significant.*

**A**

**B**

***Figure S24.*** *Me3PyC3A improves locomotor activity and Mn levels in slc39a14^-/-^ zebrafish larvae at 6 dpf.*

***A****. Boxplot of average locomotor activity of unexposed, MnCl2 (50 µM from 2 dpf), and MnCl2 & Me3PyC3A (10 µM from 2 dpf) treated larvae at 6 dpf (n=24 per group). One-Way ANOVA with Tukey's multiple comparisons test. *P=0.0116* ***B****. Mn levels determined by ICP-MS from pools of 10 larvae at 6 dpf exposed to MnCl2 (50 µM from 2 dpf) following cardiac injection at 4 and 5 dpf with Me3PyC3A (100 pg). Student’s t-test. **P= 0.0062.*

***Figure S25****. Metal-PyC3A complexes measured by LC-MS/MS from dried blood spots after* ***A, B*** *iv administration of H3PyC3A (20mg/kg) in wild-type (beige triangles) and Slc30a10^KO/KO^ mice (pink squares) and* ***C, D*** *po administration of HCL-PyC3A (80mg/kg) in Slc30a10^KO/KO^ mice (n=3 per timepoint). One Way Anova. **P<0.01, ns=not significant.*

##### References

1. Gale, E.M., Atanasova, I.P., Blasi, F., Ay, I., and Caravan, P. (2015). A Manganese Alternative to Gadolinium for MRI Contrast. Journal of the American Chemical Society *137*, 15548-15557. 10.1021/jacs.5b10748.
2. Islam, M.K., Kim, S., Kim, H.K., Park, S., Lee, G.H., Kang, H.J., Jung, J.C., Park, J.S., Kim, T.J., and Chang, Y. (2017). Manganese Complex of Ethylenediaminetetraacetic Acid (EDTA)-Benzothiazole Aniline (BTA) Conjugate as a Potential Liver-Targeting MRI Contrast Agent. J Med Chem *60*, 2993-3001. 10.1021/acs.jmedchem.6b01799.
3. Piccinelli, F., De Rosa, C., Melchior, A., Faura, G., Tolazzi, M., and Bettinelli, M. (2019). Eu(iii) and Tb(iii) complexes of 6-fold coordinating ligands showing high affinity for the hydrogen carbonate ion: a spectroscopic and thermodynamic study. Dalton Trans *48*, 1202-1216. 10.1039/c8dt03621g.
4. Coles, S.J., and Gale, P.A. (2012). Changing and challenging times for service crystallography. Chemical Science. The Royal Society of Chemistry.
